# Identification of a human Anxa1-associated dopaminergic population linked to Parkinson’s disease vulnerability and sensorimotor putaminal networks

**DOI:** 10.64898/2026.11.20.766782

**Authors:** Elan Schonfeld, Marwan N. Baliki, Mariana H.G. Monje, Behzad Yaghmaeian Salmani, Thomas Perlmann, Thomas S. Hnasko, Jean-Francois Poulin, A. Vania Apkarian, D. James Surmeier, Daniel Dombeck, Rajeshwar Awatramani

## Abstract

The selective loss of dopaminergic (DA) neurons drives motor symptoms of Parkinson’s disease (PD). In mice, *Anxa1*^+^ SNc DA neurons are highly vulnerable in PD models, project to the dorsolateral striatum, and display movement-correlated activity, yet they lack clearly defined human counterparts. We transfer the mouse taxonomy to human using a hierarchical mixture-of-experts model, recovering a human Anxa1-associated population. We find that this was the most depleted subtype in PD brains. A ligand-receptor algorithm, validated against mouse tracing, infers that these neurons preferentially innervate the caudal putamen. There, DaT-SPECT shows the earliest and most severe DA axon loss in PD. Functional connectivity positions this region as a critical node within the somato-cognitive action network, linking DA neuron vulnerability to circuit dysfunction. To translate these insights, we present an interactive portal unifying human datasets with a cross-species FDA Drug Atlas, nominating *CACNA1G* as a candidate neuroprotective target for the Anxa1-associated population.

**Highlights:**

- Transferring the mouse subtype organization to human identifies an Anxa1-associated population more depleted in PD than any existing human subtype
- The inferred preferential target of this population is the caudal putamen in humans
- The Anxa1-associated striatal field loses DAT earliest and most severely in PD and is preferentially connected to the recently identified somato-cognitive action network
- Prodromal risk localizes to this Anxa1-associated field within the posterior putamen
- DopaBase Human unifies human DA datasets with disease risk and drug-target evidence across 11 DA-related diseases
- The FDA Drug Target Atlas provides a first-pass screen for drugs targeting the DA system
- Subtype-resolved pharmacology nominates *CACNA1G* as a candidate neuroprotective target for preclinical testing in PD

## Introduction

Midbrain dopaminergic (DA) neurons regulate movement, reinforcement learning, motivation, and cognitive control – functions distributed across nigrostriatal, mesolimbic, and mesocortical projections.^1–9^ This breadth is matched by the range of disorders in which DA signaling is dysfunctional: degeneration of nigrostriatal neurons results in the hallmark motor deficits of Parkinson’s disease (PD), whereas altered transmission through dopaminergic circuitry contributes to schizophrenia, addiction, and attention-deficit/hyperactivity disorder (ADHD).^10–15^ Functions are not uniformly distributed across the DA system, as molecularly defined subpopulations display distinct vulnerabilities to disease and emerging evidence indicates that they have dissociable roles in behavior.^1–4,15–21^

A fundamental division of midbrain DA neurons separates the *Sox6*^+^ population occupying the ventral tier of the substantia nigra pars compacta (SNc) from their *Calb1*^+^ counterparts in the dorsal tier and the ventral tegmental area (VTA).^1,2^ In PD, these *Sox6*^+^ neurons degenerate to a far greater extent than *Calb1*^+^ neurons, which are relatively spared.^2,22–24^ Within this broad partition, there exists more granular variation: recent work has divided transcriptomic space into numerous distinct “territories” or “subtypes” of mouse midbrain DA neurons.^1,25–28^ Although many of these represent a continuum within transcriptomic space, these subtypes are not merely transcriptomic subdivisions – molecular identity determines projection targets, and nigrostriatal projections are topographically organized such that distinct DA populations innervate distinct striatal territories.^2,3,19–21,29,30^ Subtypes also carry distinct functional signals, some tracking movement parameters, others reward and motivation.^3,15,31–34^

Despite this organization, existing pharmacology does not resolve targets at this resolution. DA-targeted drugs can be highly selective for individual receptors or transporters, but receptor expression does not align with DA population identity – the same receptor is often expressed by cell types with unrelated functions, so a receptor-selective drug is still circuit-indiscriminate.^35–37^ Levodopa and D2 agonists restore motor function in PD at the cost of dyskinesia and impulse-control disorders, while D2 antagonists reduce psychotic symptoms in schizophrenia but also block D2 receptors in the motor striatum and pituitary, causing drug-induced parkinsonism and elevated prolactin.^38–45^ A drug confined to target DA populations that produce a specific symptom would circumvent these liabilities, but in humans, drug targets have not been mapped onto specific DA populations, and resolving disease-implicated populations remains an open challenge.^4,46,47^

The marker Annexin A1 (*Anxa1*), identified by single-cell studies has been used to demarcate a subpopulation of *Sox6*^+^ cells concentrated in the ventral SNc, which project to the dorsolateral striatum in mouse.^1,3,16,48^ *Anxa1*^+^ neurons encode movement parameters such as acceleration rather than the reward, value, or motivational signals classically attributed to DA neurons.^1,3,16,49^ In mouse models of PD, they degenerate first and most severely, and silencing them induces early motor phenotypes such as impaired movements and tremor-like symptoms.^16,48^ Several observations suggest that a related population may exist in humans, although direct proof is lacking. The mouse dorsolateral striatum is thought to correspond in primate to the posterior putamen, as both are defined by convergent input from the sensorimotor cortex; this territory is where DA loss is first and most severe in PD, whereas the caudate remains less denervated.^50–56^ Furthermore, dual in situ hybridization (ISH) for *TH* and *ANXA1* in postmortem human midbrain localizes *ANXA1*-expressing DA neurons to the ventral SNc and shows their degeneration in PD.^49^ However, these observations do not establish that the labeled cells constitute the same molecular subtype as defined in mouse. Mouse *Anxa1*^+^ DA neurons display a combinatorial gene expression profile nested within the *Sox6*^+^ family branch, whereas probe-based RNAScope in human identifies cells expressing individual transcripts and therefore cannot by itself define an equivalent molecular subtype in humans.^1,3^ Conversely, single-cell sequencing can resolve molecularly defined human DA populations, but no clear counterpart to mouse *Anxa1*^+^ DA neurons has emerged from human midbrain single-cell data.^1–4,46,57^ Diffusion MRI provides complementary circuit-level evidence, identifying a macroscale lateral SNc–posterior motor putamen–somatomotor axis consistent with primate tracing, but cannot assign this organization to a molecularly defined DA population.^58–60^ Establishing such a correspondence would require genetic access to selectively label a molecular DA subtype and trace its projections or record its activity – the approaches that enabled molecular identity, circuits, and function to be unified in mouse but cannot be applied in living humans. Overall, the field lacks an understanding of the human Anxa1 subtype, if it exists, and more generally, the correspondence between all identified mouse and human subtypes.

Three gaps therefore separate the mouse *Anxa1*^+^ population from a circuit-level account of its human counterpart. First, while *ANXA1*^+^ DA neurons have been identified in humans by in-situ hybridization^49^, single nuclei transcriptomics have thus far failed to clearly establish a corresponding population. We train and benchmark a hierarchical mixture-of-experts (HMoE) model that transfers the mouse DA hierarchy across RNA-sequencing datasets while resisting cross-species expression bias.^61,62^ Our open-source, open-weight model achieves state-of-the-art performance compared to 13 existing approaches, and identifies an Anxa1-associated population in human. We find that this population is the most depleted in PD.^63–66^ Second, subtype identity alone does not resolve projection or circuit organization – and human experimentation cannot supply it either: as a proxy, we use ligand-receptor affinity analysis on multiple independent datasets to infer subtype-specific DA communication affinity, and demonstrate that this method accurately captures canonical projection patterns in mouse.^3^ Because DA subtypes innervate the striatum topographically, these communication affinity programs define a spatial hypothesis that can be tested against regional presynaptic DAT loss measured by DaT-SPECT. We find that the inferred Anxa1-associated striatal field is preferentially depleted in early disease, captures the putaminal territory in which prodromal risk is concentrated, and lies within the sensorimotor domain of the putamen connected to the somato-cognitive action network (SCAN)^67^, as resolved by resting-state fMRI.^68–70^ Third, vulnerable populations have not been transcriptionally mapped to the pharmacological targets through which they can be manipulated. We connect disease-associated DA populations to FDA-regulated targets across eleven DA-related diseases and three species, and we deploy these analyses in DopaBase Human, an interactive, open-access atlas comprising human DA transcriptomics, disease risk, and the FDA Drug Target Atlas. Using this, we identify the CaV3.1 channel gene *CACNA1G* as a candidate target, enriched in the Anxa1-associated population, for preclinical testing in PD models. Overall, this work unifies molecular subtype identity and functional network organization, enabling us to identify and characterize a highly vulnerable Anxa1-associated human DA population.

## Results

### Hierarchical mixture-of-experts modeling resolves midbrain dopaminergic subtype organization

To preserve correspondence with previously established hierarchy of mouse DA neurons, we train a hierarchical mixture-of-experts model (HMoE) that classifies cells through a series of tiered decisions, with each node corresponding to a branch point in the Gaertner and Oram et al. reference atlas^1^. Unlike a flat multiclass classifier that assigns among all terminal subtypes simultaneously, HMoE progressively routes cells down the model’s tree-structured architecture from broad molecular families to terminal DA subtypes (**Figure 1a**).^61,62,71,72^ This reference-anchored design maps cells from new datasets onto the existing mouse subtype hierarchy^1^, rather than requiring de novo clusters to be subsequently reconciled with the reference

**Figure 1.**
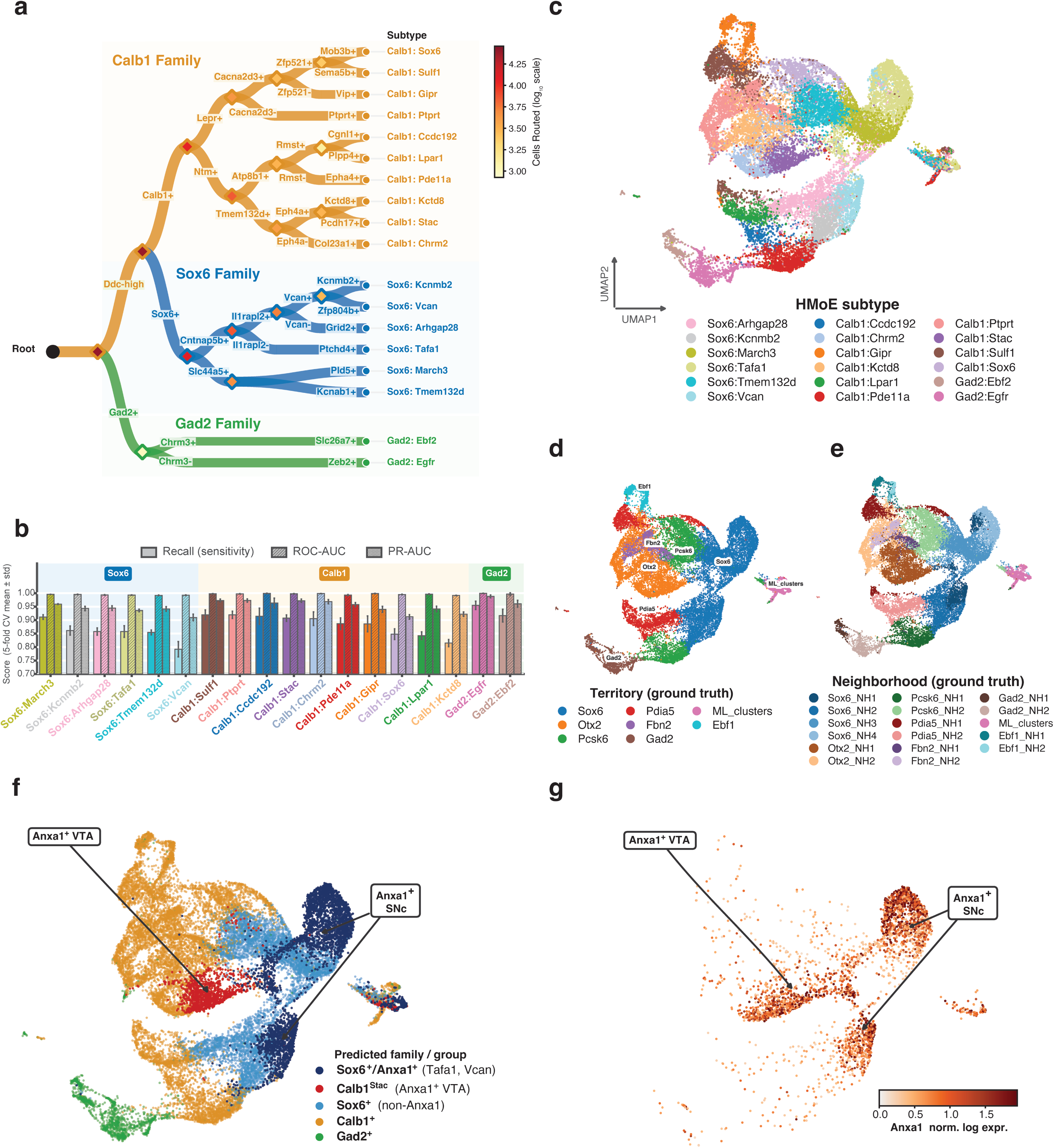
A hierarchical mixture-of-experts model resolves mouse midbrain dopaminergic subtypes and transfers to an independent atlas. **(a)** Architecture of the hierarchical mixture-of-experts (HMoE) classifier over the Gaertner–Oram et al. mouse dopaminergic taxonomy, comprising 18 terminal subtypes within the Calb1, Sox6, and Gad2 families. Diamonds represent internal binary experts and circles represent terminal subtypes. Branch labels identify marker genes defining each split; branch width and diamond color scale with the number of reference cells supported at each node. Conditional branch probabilities are propagated from the root to the terminal leaves, and the leaf with the highest probability determines subtype assignment. **(b)** Per-subtype recall (sensitivity), one-versus-rest area under the receiver operating characteristic curve (ROC-AUC), and one-versus-rest area under the precision–recall curve (PR-AUC) from five-fold stratified cross-validation, with one out-of-fold prediction per reference cell (n=24,419). Bars are ordered by recall within family; error bars indicate s.d. across folds. **(c–g)** Transfer of the frozen classifier to 24,094 author-annotated DA nuclei from six mice in the Yaghmaeian Salmani et al. dataset, comprising 18,884 nuclei from intact hemispheres and 5,210 from 6-OHDA-lesioned hemispheres. **(c)** HMoE-predicted terminal subtype. **(d)** Source-study territory annotation. **(e)** Source-study neighborhood annotation. **(f)** Predictions collapsed into *Anxa1*^+^ SNc (Sox6^Tafa1^ and Sox6^Vcan^), the *Anxa1*^+^ VTA (Calb1^Stac^) population, remaining *Anxa1*^−^/*Sox6*^+^ DA neurons, remaining *Anxa1*^−^/*Calb1*^+^ neurons, and *Gad2*^+^ neurons. Arrows mark the Anxa1-expressing VTA and SNc compartments. **(g)** Normalized Anxa1 expression on the same embedding.

In five-fold cross-validation of the model’s performance on classifying mouse DA neurons (n = 24,419), whose subtype labels were defined in the Gaertner and Oram et al. reference atlas independently of this model^1^, HMoE achieved 95.6 ± 0.4% accuracy on cells not seen during model training in distinguishing the three main DA families (**Supplemental Data Table 1**). The model was also highly capable of predicting terminal DA subtypes, achieving an accuracy of 87.4 ± 0.4% and AUROC of 0.9957 ± 0.0003 (**Figure 1b; Supplemental Data Table 1**). As additional validation, performance generalized to a 10% held-out test set of 2,442 mouse DA nuclei from the Gaertner and Oram et al. dataset, with family-level accuracy of 95.4% and subtype-level accuracy of 88.1%. (**Supplemental Data Table 1**). We next tested the model’s generalization and biological specificity, applying HMoE to a held-out snRNA-seq dataset generated independently,^25^ of 24,094 midbrain DA nuclei that was excluded from atlas construction and model training. Specifically, we asked whether HMoE can distinguish two *Anxa1*^+^ DA populations with distinct projection targets: *Sox6*^+^ SNc neurons (Sox6^Tafa1^ and Sox6^Vcan^) that project to the dorsolateral caudoputamen vs. the *Anxa1*^+^/*Otx2*^+^ VTA neurons (Calb1^Stac^) projecting to the medial shell of the nucleus accumbens.^1,30^ HMoE recovers Sox6/Calb1-family organization (**Figure 1c-e; Supplemental Figure 1a and b**) and assigned SNc and VTA *Anxa1*^+^ cells to their expected atlas subtypes (**Figure 1f and g**), with the predicted Sox6^Tafa1^, Sox6^Vcan^, and Calb1^Stac^ subtypes exhibited the highest mean *Anxa1* expression among all predicted subtypes, with *Anxa1* detected in 53%, 43%, and 40% of assigned nuclei.^25^ The separation of these predicted SNc and VTA populations despite their shared *Anxa1* expression is consistent with predictions reflecting broader transcriptional programs rather than individual markers alone.

### HMoE recovers human dopaminergic family structure and identifies an Anxa1-associated population

We next asked whether HMoE generates biologically meaningful categorization of human midbrain DA neurons. To test this hypothesis, the model was applied to 22,048 human SNc DA neurons and benchmarked against independently annotated Sox6/Calb1 family labels in the 20,630 human DA neurons from Kamath and Abdulraouf et al. (**Figure 2a**; Gad2 nuclei excluded for quality).^4^ Despite the mouse-human transcriptional divergence, the HMoE model remained robust, achieving an accuracy of 81.7% (80.5% when Gad2-predicted cells are reclassified rather than excluded) and outperforming 13 baseline methods (**Figure 2b and 2c; Supplemental Figure 2e**) for label transfer, including traditional machine learning (ML) algorithms, scVI/scANVI integration models, SingleCellNet, and the scGPT foundation model.^63–66^ Unlike these general-purpose approaches, HMoE is structured around the biological hierarchy of the classification problem, resolving broad dopaminergic families before progressively finer identities. In an independent dataset of 823 midbrain DA neurons from Siletti et al.^46^, HMoE also recovered Sox6-, Calb1-, and Gad2-like organization, supporting generalization beyond the Kamath dataset (**Supplemental Figure 2b**).

**Figure 2.**
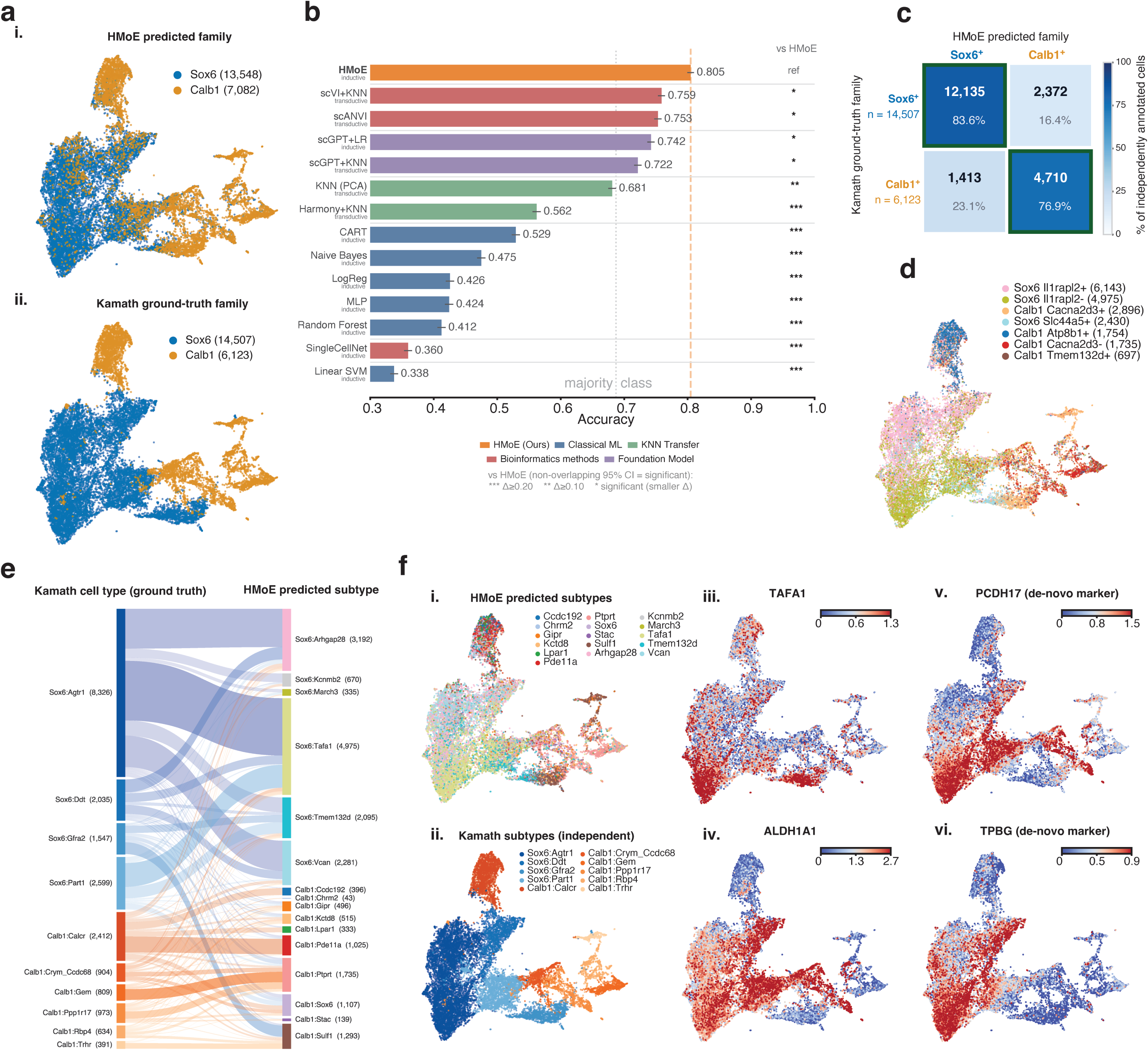
The mouse-trained HMoE transfers to human dopaminergic neurons and resolves an Anxa1-associated population. **(a)** HMoE-predicted Sox6/Calb1 family identity (i) and independently annotated Kamath family identity (ii) on the same UMAP after exclusion of 1,418 Gad2-like profiles (n=20,630 nuclei). **(b)** Cross-species label-transfer accuracy for HMoE and 13 baseline methods evaluated across all 22,048 Kamath DA nuclei. Outputs from every method were reduced to normalized for assignment to either Sox6 and Calb1 families, this assigns Gad2-like profiles according to their relative Sox6/Calb1 support. The dashed line marks majority-class accuracy, and labels beneath method names indicate inductive or transductive transfer. All baseline Wilson 95% confidence intervals lay below the HMoE interval. Asterisks encode the absolute accuracy difference from HMoE rather than p-value thresholds: * Δ < 0.10, ** 0.10 ≤ Δ <0.20, and *** Δ ≥ 0.20. **(c)** Family-level confusion matrix after exclusion of the Gad2-like profiles (n=20,630). Fill indicates row recall; cells report counts and row percentages, and green outlines identify the dominant prediction. Overall accuracy was 81.7%. **(d)** The same nuclei colored by assignments at the height-5 level of the HMoE hierarchy from Fig. 1a, representing an intermediate level between broad families and terminal subtypes. **(e)** Sankey diagram relating the ten Kamath cell-type annotations to the 16 HMoE terminal subtypes predicted in human. Each box represents a DA cell type or subtype, and its height is proportional to the number of cells assigned to it. **(f)** HMoE-predicted subtype showing an Anxa1-associated subtype in light green (i), Kamath cell type (ii), and ALRA-imputed expression of *TAFA1* (iii), *ALDH1A1* (iv), *PCDH17* (v), and *TPBG* (vi) on the Kamath UMAP, showing enriched expression in the Anxa1-associated subtype.

At dendrogram height 5, which partitions the *SOX6*^+^ and *CALB1*^+^ families into seven higher-order branch groups (3 *SOX6*^+^, 4 *CALB1*^+^), cells assigned to each branch formed clean, sharply delineated regions of the UMAP embedding (**Figure 2d**), whereas at terminal-subtype resolution, assignments became more intermingled within these broader regions (**Figure 2e and f**). HMoE assigned Kamath nuclei to 16 terminal subtypes – six Sox6 (Tafa1, Vcan, Arhgap28, March3, Tmem132d, Kcnmb2) and ten Calb1 (Ccdc192, Chrm2, Gipr, Kctd8, Lpar1, Pde11a, Ptprt, Sox6, Stac, Sulf1) – with no nuclei retained in the Gad2 family after the exclusion described above (**Figure 2e**). We then focused on cells assigned to the subtypes associated with the vulnerable mouse *Anxa1*^+^ SNc: Sox6^Tafa1^ and Sox6^Vcan^; we found that these were predicted with median confidence = 0.998 and 0.940, respectively (**Supplemental Figure 2c**; hereafter termed Anxa1-associated). At this finer resolution, 7,256 DA neurons were assigned to Anxa1-associated subtypes, the most frequent Kamath et al. subtypes were SOX6_AGTR1 (61.9%), SOX6_PART1 (17.1%), and SOX6_DDT (11.9%), placing the predicted population predominantly within the ventral-tier-enriched SOX6_AGTR1 subtype (**Figure 2e**).^4^ Other markers with enriched detection and expression in this predicted subset included *PCDH17* (LFC = +0.91; FDR < 0.001), consistent with mouse expression patterns^25^, and *TPBG* (LFC = +1.23; FDR < 0.001) (**Figure 2f; Supplemental Figure 2d**). Despite scANVI achieving broad-family transfer accuracy of 0.75, comparable to HMoE, the methods diverged substantially at terminal-subtype resolution: scANVI assigned only 100 cells to Sox6^Tafa1^ and redistributed much of the corresponding compartment to Sox6^Arhgap^^28^, whereas HMoE retained 4,975 Sox6^Tafa1^ assignments (**Supplemental Figure 2e**). Thus, methods with competitive broad-family transfer could nonetheless predict negligible numbers of Anxa1-associated cells, collapsing a biologically critical branch relevant to PD. Overall, HMoE identified a subset of human DA neurons distributed across multiple previously defined transcriptomic clusters as analogous to the mouse *Anxa1*^+^ cells most vulnerable in PD.

### Anxa1-associated DA neurons exhibit heightened vulnerability to PD

One validation of predicted Anxa1-associated cells would be to examine their heightened vulnerability as demonstrated in mouse studies.^16,48^ We therefore asked 1. whether Anxa1-associated SNc DA neurons are preferentially lost in PD and 2. whether they carry elevated common-variant genetic risk. First, we examined the depletion of this population directly. In paired hemispheric analyses of a mouse 6-OHDA model snRNA-seq dataset^25^, with Calb1 cells as reference, preservation declined from Calb1 to Sox6 non-Anxa1 (log_2_OR = –0.64) and Calb1 vs. Anxa1-associated (log_2_OR = –0.80, p vs. Sox6 = 0.258) (**Figure 3b**). To test whether this same comparison holds in postmortem human PD midbrain with documented pathological midbrain DA neuron loss, we performed MASC analysis to compare control donors (n = 8) to PD donors (n = 7, Lewy body dementia brains were excluded here), controlling for sex, age, and post-mortem interval.^4^ Relative to Calb1, the SOX6_AGTR1 cluster was depleted (log_2_OR = –1.81), reproducing the subtype previously reported as the most vulnerable population in PD.^4^ The Sox6 family as a whole, however, was depleted to the same degree (log_2_OR = –1.85), and tested directly against the remainder of the family, SOX6_AGTR1 showed no excess depletion (p = 0.598) – its vulnerability is a property of the Sox6 family rather than of the subtype. Strikingly, Anxa1-associated neurons were more aggressively depleted than Calb1 (log2OR = –2.77) and than both non-Anxa1 SOX6_AGTR1 (p = 0.010) and non-Anxa1 Sox6 (p = 0.002) (**Figure 3c**). Consistent with the Anxa1-associated population being responsible for vulnerability within these broader groups, its removal shifted SOX6_AGTR1 depletion from log_2_OR = –1.81 to –0.93 and Sox6 depletion from –1.85 to –1.11 (**Figure 3c**). Analysis of cell density on the UMAP plot concordantly localized the Anxa1-associated manifold to the disease-depleted end of a control vs. PD depletion gradient (**Figure 3d**). The Anxa1-associated group was depleted in PD, in Lewy body dementia (LBD; n=3), and in the pooled PD + LBD comparison (FDR < 0.01 in all three) (**Supplemental Figure 3a–f**). The effect did not depend on the subtype classification method, appearing equivalently in scANVI predictions (**Supplemental Figure 3g**), and was absent in size-matched random populations drawn from either the Sox6 family or the SOX6_AGTR1 subtype, confirming that the excess depletion is specific to the Anxa1-associated population rather than to any subpopulation of comparable number of cells (**Figure 3e**).

**Figure 3.**
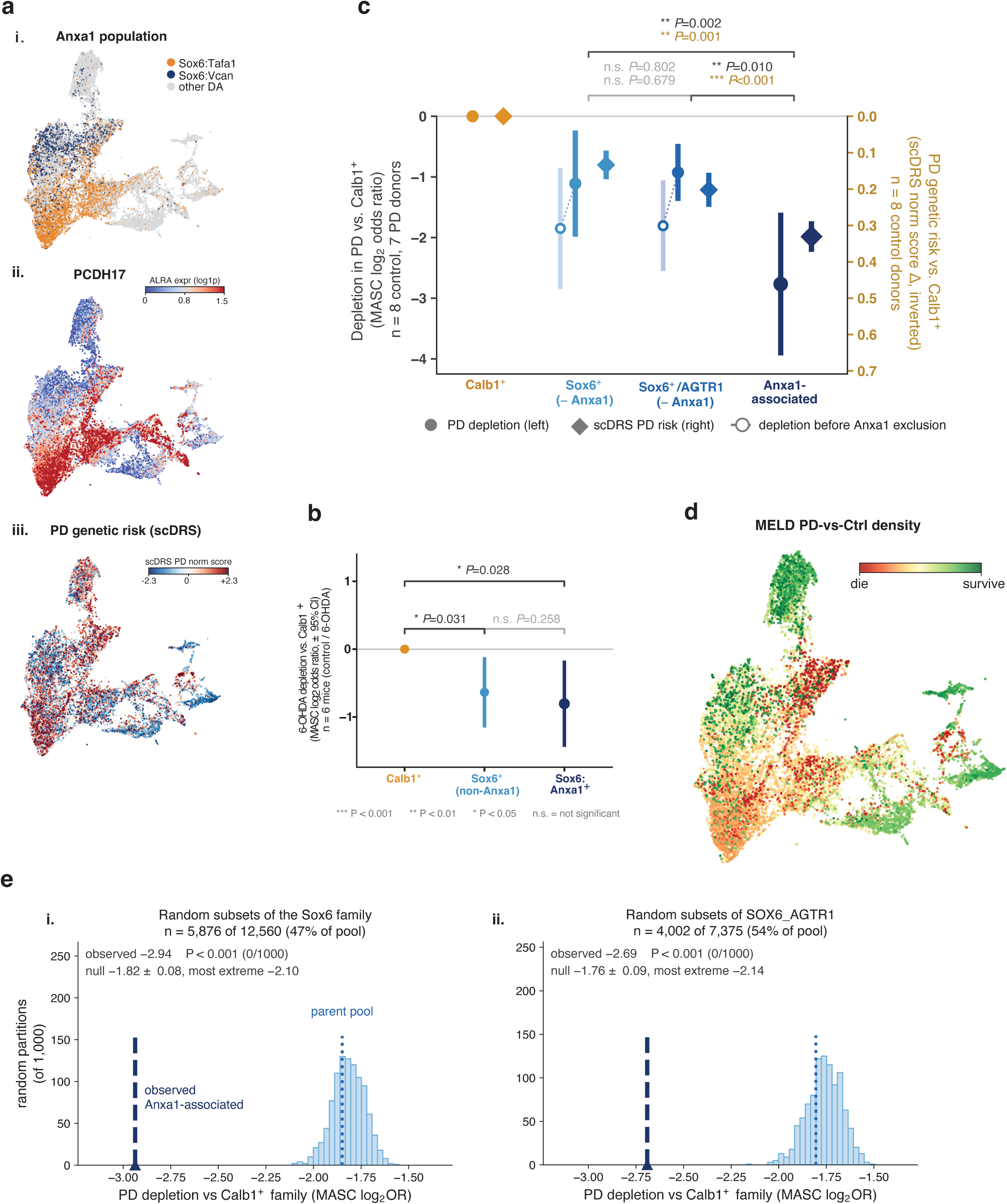
Anxa1-associated human dopaminergic neurons are selectively depleted in Parkinson’s disease and carry elevated polygenic risk. **(a)** Kamath DA-neuron UMAP showing HMoE-defined Anxa1-associated cells – Sox6^Tafa1^ and Sox6^Vcan^ (i) – ALRA-imputed *PCDH17* expression (ii), and normalized PD scDRS scores (iii). **(b)** Relative population abundance following unilateral 6-OHDA lesion in mouse, estimated by MASC generalized linear mixed models using Calb1 neurons as the reference (n=6 mice with paired intact and lesioned hemispheres). Negative log_2_ odds ratios indicate relative depletion. Whiskers show Wald 95% confidence intervals. The two brackets against Calb1 restate the corresponding model estimates; the Anxa1 versus non-Anxa1 Sox6 bracket is a paired t-test across the six animals on within-animal log2 abundance ratios. **(c)** Human PD depletion and PD polygenic-risk enrichment for Calb1, remaining Sox6, remaining SOX6_AGTR1, and Anxa1-associated neurons in eight control and seven PD donors. Anxa1-associated cells were removed from both Sox6 comparator populations. Circles and the left axis show MASC log2 odds ratios adjusted for sex, age, and postmortem interval in eight control and seven PD donors; open circles show the same populations before the Anxa1-associated cells were removed. Diamonds and the inverted right axis show scDRS effects relative to Calb1, estimated in the eight control donors by a linear mixed model with a donor random intercept. Whiskers indicate Wald 95% confidence intervals. Brackets show donor-level pairwise contrasts: within-donor log2 abundance ratios for depletion, and random-effects meta-analysis of within-donor score differences for scDRS. **(d)** MELD relative PD-versus-control density on the same UMAP. Red indicates local enrichment in PD and green indicates enrichment in controls; MELD is shown descriptively, whereas population-level inference was performed using MASC. **(e)** Empirical null distributions generated from 1,000 size-matched random subsets of the Sox6 family (i) and SOX6_AGTR1 population (ii). Dashed lines mark the observed depletion of the Anxa1-associated population; P-values are one-sided empirical p-values. In b and c, *p < 0.05, **p < 0.01, and ***p < 0.001; n.s., not significant.

A second validation of heightened vulnerability would be preferential expression of PD GWAS-implicated genes in this population. Single-cell disease relevance scores computed from the Nalls et al. PD GWAS confirmed heightened genetic vulnerability of Anxa1-associated neurons to PD (Δ = +0.33) over both the previously identified vulnerable SOX6_AGTR1 subtype (Δ = +0.20; p < 0.001) and the *SOX6*^+^ family (Δ = +0.13; p = 0.001) (**Figure 3a and c**).^73,74^ Together, these findings identify a human DA population, aligned with the mouse *Anxa1*^+^ SNc, that is more severely depleted in PD than any previously described subtype and carries elevated PD genetic risk.

### Communication affinity analysis predicts an Anxa1-associated caudal putaminal territory

To predict the striatal targets of midbrain DA neuron subtypes, we applied a communication affinity analysis using curated ligand–receptor pairs from the LIANA consensus resource.^75,76^ For each DA subtype–striatal region combination, we scored communication potential by combining ligand expression in the DA subtype with expression of the corresponding receptors in regional SPNs. We then used a Tensor-cell2cell–based decomposition to identify shared patterns across ligand–receptor pairs, DA subtypes and striatal regions, from which we reconstructed each subtype’s relative regional target-affinity map (**Methods**).^76^ Because this approach infers communication potential rather than measuring axonal connectivity, we first tested it in mouse, recovering the canonical dorsolateral-to-ventromedial gradient from *Anxa1*^+^ → *Sox6*^+^ → *Calb1*^+^: striatal hotspot centroids shift progressively from ventrolateral to dorsomedial (0.63 → 0.56 → 0.52 mm) across the three groups (p = 0.018) (**Figure 4a and b**).^2,3^ Across two independent datasets it also assigns the *Sox6*^+^ family to the caudoputamen (p < 0.001) and the *Calb1*^+^ family to the nucleus accumbens (p = 0.01) and olfactory tubercle (p < 0.001) (**Figure 4a; Supplemental Figure 4a**).^2,3^ We then compared the predictions to subtype-specific experimental viral tracings^30^ (**Figure 4c**), finding concordance across top projection planes corresponding to each of *Anxa1*^+^ (AUC =0.91), *Sox6*^+^ (AUC = 0.75), and *Calb1*^+^ (AUC = 0.92) families, as well as rostral-to-caudal gradient agreement between the two methods.

**Figure 4.**
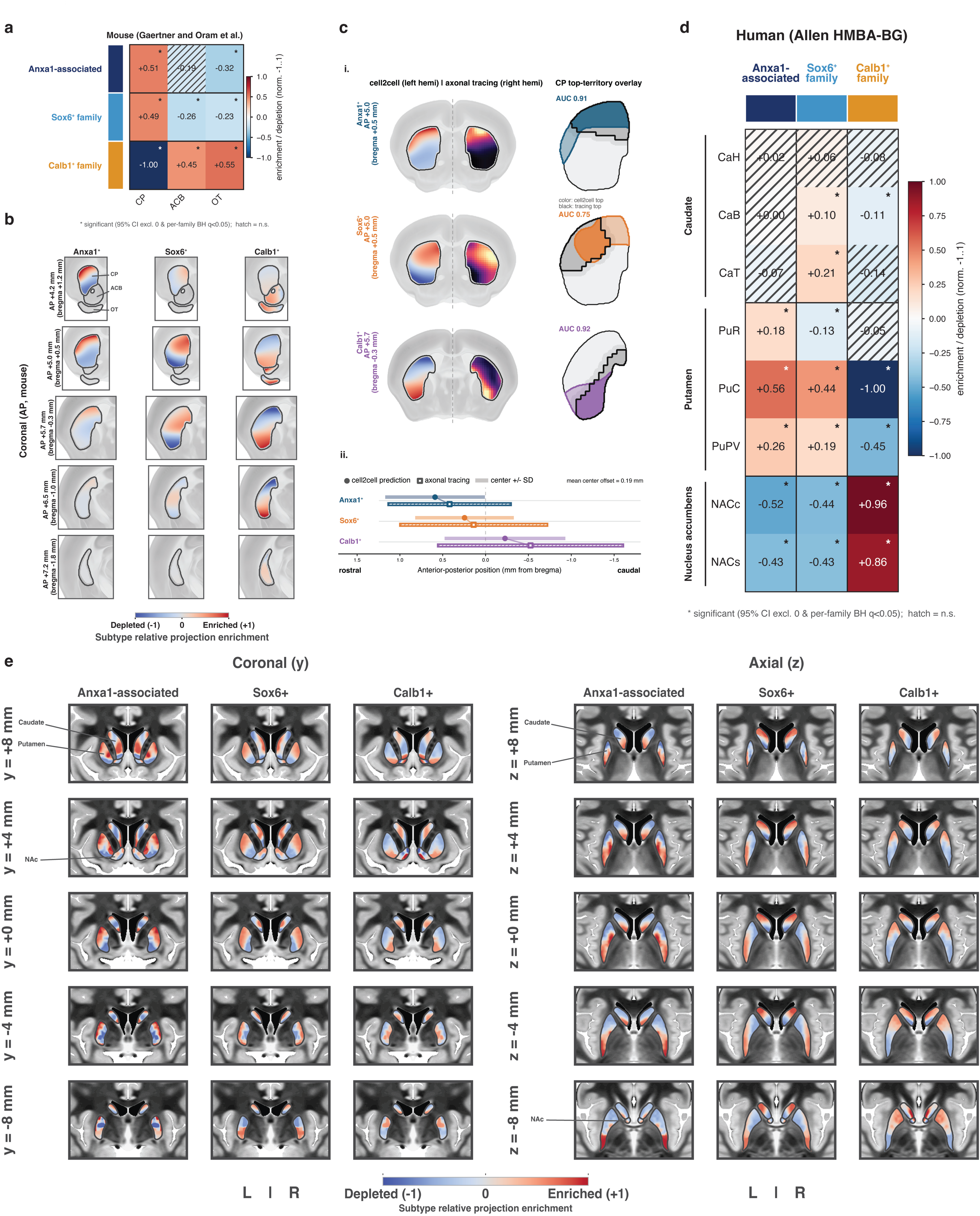
Mouse tracing validates communication affinity-derived maps of subtype-specific striatal target affinity across species. Heatmaps show relative communication-affinity enrichment for three DA sender groups: Anxa1-associated, Sox6-family, and Calb1-family neurons. Values are scaled from −1 to +1 within each analysis; positive values indicate greater communication affinity than the other DA groups for that receiver region. Asterisks identify cells whose bootstrap 95% confidence interval excludes zero and whose within-family Benjamini–Hochberg-adjusted q-value is <0.05; hatching denotes cells failing either criterion. **(a)** Mouse communication affinity across caudoputamen (CP), nucleus accumbens (ACB), and olfactory tubercle (OT), inferred from Gaertner–Oram DA senders and spatially resolved striatal receivers. **(b)** Anatomical rendering of the same mouse communication affinity fields across five coronal levels. Color indicates enrichment or depletion relative to the other DA groups. **(c)** Validation against published subtype-specific anterograde axonal tracing. Communication affinity-derived and tracing maps are displayed on opposite hemispheres at the coronal level of peak projection for each group (i), with overlays comparing their highest-affinity territories. ROC-AUC values were 0.91 for Anxa1-associated, 0.75 for Sox6-family, and 0.92 for Calb1-family maps. Anteroposterior projection centers are compared in (ii); the mean center offset was 0.19 mm, and the rostrocaudal ordering of all three groups was preserved. **(d)** Corresponding communication-affinity map across eight human striatal regions in the Allen HMBA-BG dataset. CaH, CaB, and CaT denote caudate head, body, and tail; PuR, PuC, and PuPV denote rostral, caudal, and posterior putamen; NACc and NACs denote nucleus accumbens core and shell. **(e)** Human communication-affinity fields spatially rendered using REMAP trained jointly on the Allen HMBA-BG MERSCOPE and Kraft Slide-tags references. Coronal and axial sections are shown for each DA sender group.

Having validated our approach in mice, we next analyzed inferred communication affinity between HMoE predicted DA subtypes and human SPN datasets.^4,77,78^ We find the greatest communication affinity between the Anxa1-associated SNc population and the caudal (PuC) and posterior ventral putamen (PuPV), the subregions most affected by dopaminergic denervation of the striatum in early PD (**Figure 4d**).^4,53–55,77,79^. Conversely, this population’s communication affinity was lowest in the nucleus accumbens (**Figure 4d**). This pattern replicated in independent Slide-tags data^79^: Anxa1-associated enrichment was highest in a sensorimotor putaminal zone, while Calb1 favored a NAc-enriched zone (**Supplemental Figure 4c**). Furthermore, using a macaque dataset^77^, a similar caudal putaminal trend was observed (**Supplemental Figure 4b**). To further investigate spatial gradients between subtype-specific projections, we utilized spatial transcriptomic data of the human striatum to render the regional target-affinity estimates at a finer spatial resolution within the putamen, localizing inferred Anxa1-associated communication affinity to the dorsolateral putamen and Calb1-family communication affinity the ventromedial striatum (**Figure 4e**).^77^ Although this map was imputed, it defined an independently testable hypothesis: if Anxa1-associated vulnerability is reflected in the target field, that territory should show 1. disproportionate presynaptic dopaminergic loss in living patients with PD and 2. possible enriched connectivity to recently-defined SCAN domains, that show increased functional connectivity to various basal ganglia structures in PD.^67,80^ These tests provide an orthogonal validation of the inferred communication affinity architecture using independent in vivo imaging modalities, asking whether the predicted Anxa1-associated target field exhibits the expected dopaminergic denervation and sensorimotor cortical connectivity. The results are presented in the following sections.

### In vivo DaT depletion in Parkinson’s disease converges on the inferred Anxa1-associated putaminal territory

To determine if the inferred Anxa1-associated striatal territory is preferentially affected in PD patients, we reconstructed baseline DaT-SPECT scans from 1,127 patients with PD and 230 healthy controls drawn from the Parkinson’s Progression Markers Initiative (PPMI) dataset (**Figure 5a**).^81^ Baseline PD scans were early-stage de novo acquisitions (median 2.0 years since motor onset, interquartile range 1.1 to 3.3). Projection-weighted putaminal depletion was greatest in the Anxa1-associated field (69%) vs. the Sox6 (66%), non-Anxa1 Sox6 (46%), and Calb1 family (42%) fields (**Figure 5b**), with excess within-patient depletion (Δ) of +2 percentage points over Sox6 (rank-biserial correlation = 0.89; p < 0.001), +20 over non-Anxa1 Sox6 (rbc = 0.91; p < 0.001), and +24 over Calb1 (rbc = 0.94; p < 0.001) (**Figure 5c**). As an independent, voxelwise test, we correlated the voxelwise DaT-depletion map with each family’s projection field across the striatum. Depletion was positively concordant with the Anxa1-associated map (ρ = +0.77; spatial-null p < 0.001) and inversely concordant with Calb1 (ρ = -0.76; p < 0.001), whereas the Sox6-all (ρ = +0.15; p = 0.524) and Sox6-non-Anxa1 (ρ = -0.40; p = 0.124) maps were not significant (**Figure 5d**). This concordance was strongest in the less-affected (LA) hemisphere, where dopaminergic signal is not yet floored (**Supplemental Figure 5a**). At the early clinical stage of disease progression – <1 year since motor symptom onset (n = 255) – the Anxa1-associated territory has already reached 49% depletion, significantly greater than Sox6 (paired Δ = +8; rbc = 0.54; p < 0.001) and non-Anxa1 Sox6-associated depletion (paired Δ = +16; rbc = 0.70; p < 0.001) (**Figure 5b; Supplemental Figure 5f**).

**Figure 5.**
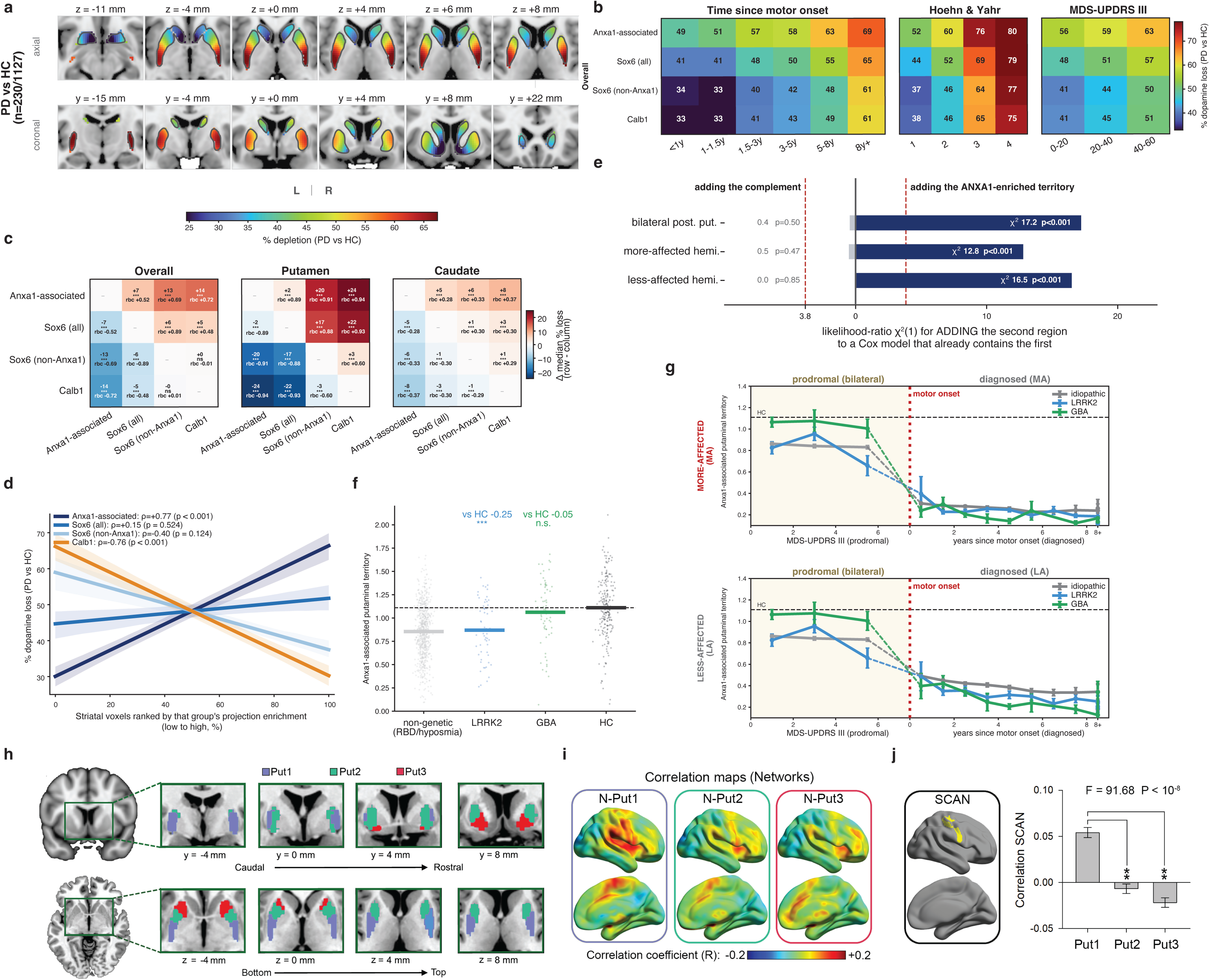
The Anxa1-associated putaminal field is linked to a sensorimotor network and exhibits early, disproportionate DAT depletion in Parkinson’s disease. **(a)** Age- and sex-adjusted percentage DAT depletion in 1,127 participants with PD relative to 230 healthy controls, shown across axial and coronal sections. **(b)** Communication affinity-field-weighted DAT depletion across time since motor onset, Hoehn and Yahr stage, and MDS-UPDRS part III severity. For each group and stage bin, the age- and sex-adjusted group depletion map was weighted by the positive portion of that group’s communication-affinity field. Values are spatial means from the corresponding group-level maps. **(c)** Within-participant differences in projection-weighted depletion among the Anxa1-associated, Sox6-all, Sox6-non-Anxa1, and Calb1 fields in the dorsal striatum, putamen, and caudate (n=1,127 baseline PD scans). Cells report the median paired difference, matched-pairs rank-biserial correlation, and Holm-adjusted P value from paired Wilcoxon signed-rank tests. **(d)** Voxelwise DAT depletion as a function of communication affinity-field enrichment across the bilateral striatum. Points represent projection-enrichment bins used for visualization; lines are linear fits and ribbons are spatial block-bootstrap 95% confidence intervals. Spearman correlations and two-sided BrainSMASH spatial-null p-values are shown. **(e)** Added prognostic information for PD phenoconversion in 1,665 prodromal participants, including 198 conversions. Bars show one-degree-of-freedom likelihood-ratio statistics from Cox models when either the Anxa1-enriched or complementary posterior-putamen field was added after the other; all models additionally included age and sex. Analyses are shown bilaterally and for the more- and less-affected hemispheres. **(f)** Prodromal posterior-putamen SBR in non-genetic participants enrolled for RBD or hyposmia, LRRK2 carriers, GBA carriers, and healthy controls with MDS-UPDRS part III ≤ 3. Points represent participants and horizontal bars indicate group means; displayed contrasts against healthy controls were adjusted for age and sex. **(g)** Anxa1-field-weighted putamen SBR across the prodromal and diagnosed disease course in idiopathic PD, LRRK2 carriers, and GBA carriers. Prodromal values are shown against MDS-UPDRS part III and diagnosed values against years since motor onset, separately for the clinically defined more- and less-affected hemispheres. Points and error bars show bin means and s.e.m.; the vertical dashed line marks motor onset. Prodromal bins are shown only where both carrier groups have at least 5 participants (MDS-UPDRS III < 7); prodromal denotes enrolment without a PD diagnosis, not absence of motor signs. Note: since the number of genetic patients tracked from prodromal to phenoconversion is low (19 LRRK2 patients and 11 GBA patients with DaTSCANS on both sides of diagnosis), patients defined as prodromal vs. de novo PD here are not necessarily the same patients. **(h)** Resting-state fMRI parcellation of the bilateral putamen into three spatial territories, Put1–Put3, shown across coronal and axial sections. **(i)** Cortical resting-state correlation maps associated with each putaminal territory, denoted N-Put1, N-Put2, and N-Put3. **(j)** Somato-cognitive action network (SCAN) reference map (yellow; left) and spatial correlation of each parcel-associated cortical correlation map with SCAN (right). N-Put1 showed the strongest positive correspondence with SCAN, whereas N-Put2 and N-Put3 showed near-zero or negative correlations (F = 91.68, p < 0.001). Brackets indicate pairwise comparisons with N-Put1 (**p < 0.01). Bars show mean ± 95% CI (n = 100).

We next asked whether genetic forms of PD show the same preferential loss of the Anxa1-associated territory, and whether that loss begins before diagnosis in non-manifest carriers. Genetic comparisons drew on three PPMI cohorts imaged under one DaT-SPECT protocol: diagnosed PD (937 idiopathic, 124 LRRK2, 63 GBA participants), the Prodromal cohort (non-carriers enrolled for RBD or hyposmia, plus LRRK2 and GBA carriers without a PD diagnosis), and healthy controls. Because only 5 LRRK2 and 3 GBA prodromal carriers phenoconverted during follow-up, the prodromal and diagnosed cohorts comprise different participants, and the genotype trajectories reported below are cross-sectional. GBA-PD additionally showed excess putaminal loss (**Supplemental Figure 5e**). To separate genotype from differences in disease duration and severity between cohorts, we fitted sequential models across all visits. The LRRK2 difference attenuated once disease time was accounted for, whereas the GBA deficit persisted through sequential adjustment for age, sex, time since motor onset, and Hoehn and Yahr stage, and in a disease-time–matched analysis (whole-putamen ΔSBR = –0.125, p < 0.001; –0.079 versus LRRK2, p = 0.003) (**Supplemental Figure 5d**). This excess loss localized to the Anxa1-associated putaminal territory: after adjustment for age, sex, and disease time, GBA carriers had 24% (less-affected hemisphere, p = 0.005) and 28% (more-affected, p = 0.008) lower territory DAT than idiopathic patients, versus 3% and 10% in the caudate (both n.s.), and the disease-time-matched territory deficit was detectable in the less-affected hemisphere (ΔSBR = –0.074, p = 0.024) (**Supplemental Figure 5b**). Non-manifest carriers suggested two routes to this deficit. At baseline (MDS-UPDRS III ≤ 3), LRRK2 carriers already had lower Anxa1-territory DAT than controls (ΔSBR = –0.25, p < 0.001; n = 56), whereas GBA carriers did not (ΔSBR = –0.05, p = 0.23; n = 65), consistent with the whole-putamen pattern reported by Simuni et al. and localizing it to the Anxa1-associated territory (**Figure 5f**).^82^ In both genotypes, Anxa1-territory DAT fell sharply at motor onset (GBA: 1.08 at the last prodromal bin to 0.40 in the less-affected and 0.24 in the more-affected hemisphere within the first year), after which the GBA deficit relative to idiopathic patients grew from –0.05 SBR within two years of onset (p = 0.40) to –0.15 at four to eight years (p = 0.002). This widening was confirmed by patient-level bootstrap (p = 0.048), and was consistent with the accelerated progression reported for GBA-associated PD (**Figure 5g**).^83,84^

To determine whether prodromal conversion risk was preferentially associated with the Anxa1-associated territory rather than an equivalently sized portion of the surrounding posterior putamen, we compared nested Cox models using the Anxa1-associated field and a volume-matched complementary posterior-putamen field. Adding Anxa1-associated field depletion to a model that already contained the complementary posterior-putamen field improved prediction bilaterally and in each hemisphere individually (all p < 0.001), whereas adding the complementary field after the Anxa1-associated field did not (all p ≥ 0.25) (**Figure 5e**). Thus, within this volume-matched comparison, the independent prognostic signal preferentially localized to the Anxa1-associated field.

Baseline depletion in the Anxa1- and Calb1-associated fields also predicted different clinical trajectories. For each clinical measure, we compared regression coefficients relating baseline Anxa1-versus Calb1-field depletion to subsequent clinical change. Tremor and motor measures shifted toward the Anxa1-associated side of the Anxa1–Calb1 axis, whereas neuropsychiatric and cognitive measures were shifted toward the Calb1 field (**Supplemental Figure 5c**). Because the fields’ shares of loss sum to a constant at fixed total burden, we asked whether where loss concentrates, rather than how much, carries prognostic information. For each field, excess loss was defined as the residual of field depletion after regressing out total striatal burden, age, and sex. Excess loss in the Anxa1 field predicted conversion to PD (HR = 1.18 per SD; p = 0.018), whereas excess loss in the Sox6 (HR = 0.74; p < 0.001) and Calb1 fields (HR = 0.72; p < 0.001) was associated with lower conversion, each field fitted in a separate model; 17.1% of participants with above-median Anxa1-concentrated loss converted within 3 years, versus 12.1% and 10.4% for the corresponding Sox6 and Calb1 splits (**Supplemental Figure 5c**). The same distinction separated prodromal participants enrolled for polysomnography-confirmed iRBD (n = 404) from those enrolled for hyposmia or genetic risk (n = 1,261), adjusting for total burden, age, and sex: each SD of Anxa1-field depletion raised iRBD odds 1.89-fold (p < 0.001), Sox6-field depletion 1.60-fold (p = 0.004), while Calb1-field depletion was inversely associated (OR 0.33; p < 0.001). Thus, relative Calb1-field preservation marked iRBD cases (**Supplemental Figure 5c**). Because both analyses (iRBD and phenoconversion) adjusted for total striatal burden, excess loss in one field implies relative sparing of the others. The three field estimates therefore describe a single distribution of loss across the striatum, and the inverse associations (Sox6 and Calb1 fields for conversion, the Calb1 field for iRBD) mark relative sparing of those fields.

### fMRI-defined putaminal territories link Anxa1-associated DA signaling to a sensorimotor network

To place the molecularly inferred Anxa1-associated projection field within a cortical functional network, we used resting-state fMRI from 535 healthy individuals from the NITRC 1000 Functional Connectomes Project^85^ to cluster putamen voxels according to their cortical connectivity profiles.^68–70^ The whole-brain connectivity patterns of the resulting three putaminal clusters (Put1–Put3) were then independently characterized in a separate cohort of 100 healthy participants from the Human Connectome Project (HCP)^86^, using voxelwise correlation analysis. The three clusters exhibited distinct whole-brain connectivity patterns (N-Put1–N-Put3; **Figure 5h and i**). Since increased SCAN connectivity to key basal ganglia nuclei has been causally linked to PD^80^, we asked if these putaminal clusters’ connectivity maps exhibited differential spatial correspondence with the somato-cognitive action network (SCAN).^67^ The caudal Put1 territory showed the strongest positive correspondence with SCAN, significantly exceeding N-Put2 and N-Put3, whose correlations were near zero or negative (F = 91.68, P < 0.001; **Figure 5j**). Notably, Put1 spatially overlapped the caudal putaminal field independently prioritized by ligand–receptor analysis for Anxa1-associated DA signaling (ρ = 0.80, q = 0.01) and, similarly, with the overlapping Sox6-all field (ρ = 0.79, q = 0.01), while showing inverse correspondence with Sox6-rest (ρ = −0.72, q = 0.012) and Calb1 (ρ = −0.80, q = 0.01). Put2 showed the complementary within-Sox6 pattern, correlating positively with Sox6-rest (ρ = 0.72, q = 0.01) and negatively with Anxa1-associated projections (ρ = −0.56, q = 0.019). Put3 did not reach significance for any positive association but was most aligned with the Calb1 field (ρ = 0.37, q = 0.32). Together, these results distinguish Anxa1-associated and Sox6-rest projection topographies across the connectivity-defined putaminal territories and place the Anxa1-associated field within the caudal Put1 territory, whose cortical network showed the strongest correspondence with SCAN. Thus, two independent approaches, including molecularly inferred projection mapping and resting-state functional connectivity, converged on a caudal putaminal territory associated with sensorimotor and somato-cognitive action network.

### An eleven-disease FDA Drug Target Atlas maps pharmacological access to disease-associated DA populations across species

Since molecularly defined DA populations are likely parts of distinct circuits^30^, we aimed to 1. assign disease risk to each subtype for common DA related disorders, and 2. Define FDA approved therapies that might target such vulnerable circuits, towards circuit specific therapies. First, extending our established single-cell disease relevance scoring (scDRS) pipeline^1,73^, we mapped genetic vulnerability across human, mouse, and macaque populations for ten additional diseases in which DA dysfunction is implicated: attention-deficit/hyperactivity disorder, anxiety, bipolar disorder, chronic pain, major depressive disorder, nicotine dependence, obsessive-compulsive disorder, opioid use disorder, schizophrenia, and fibromyalgia (**Figure 6d**).^1,73,74,87–96^ Disease relevance scores showed distinct subtype enrichment across these disorders (**Figure 6d**). PD GWAS signal was most strongly enriched in Sox6^Tafa1^, whereas fibromyalgia, chronic pain, and opioid use disorder were more enriched in specific Calb1 subtypes, such as Calb1^Sulf1^. These analyses point to DA subcircuits that may be affected in each disorder. This is particularly valuable for disorders such as these, many of which lack well-defined cellular substrates or circuits.

**Figure 6.**
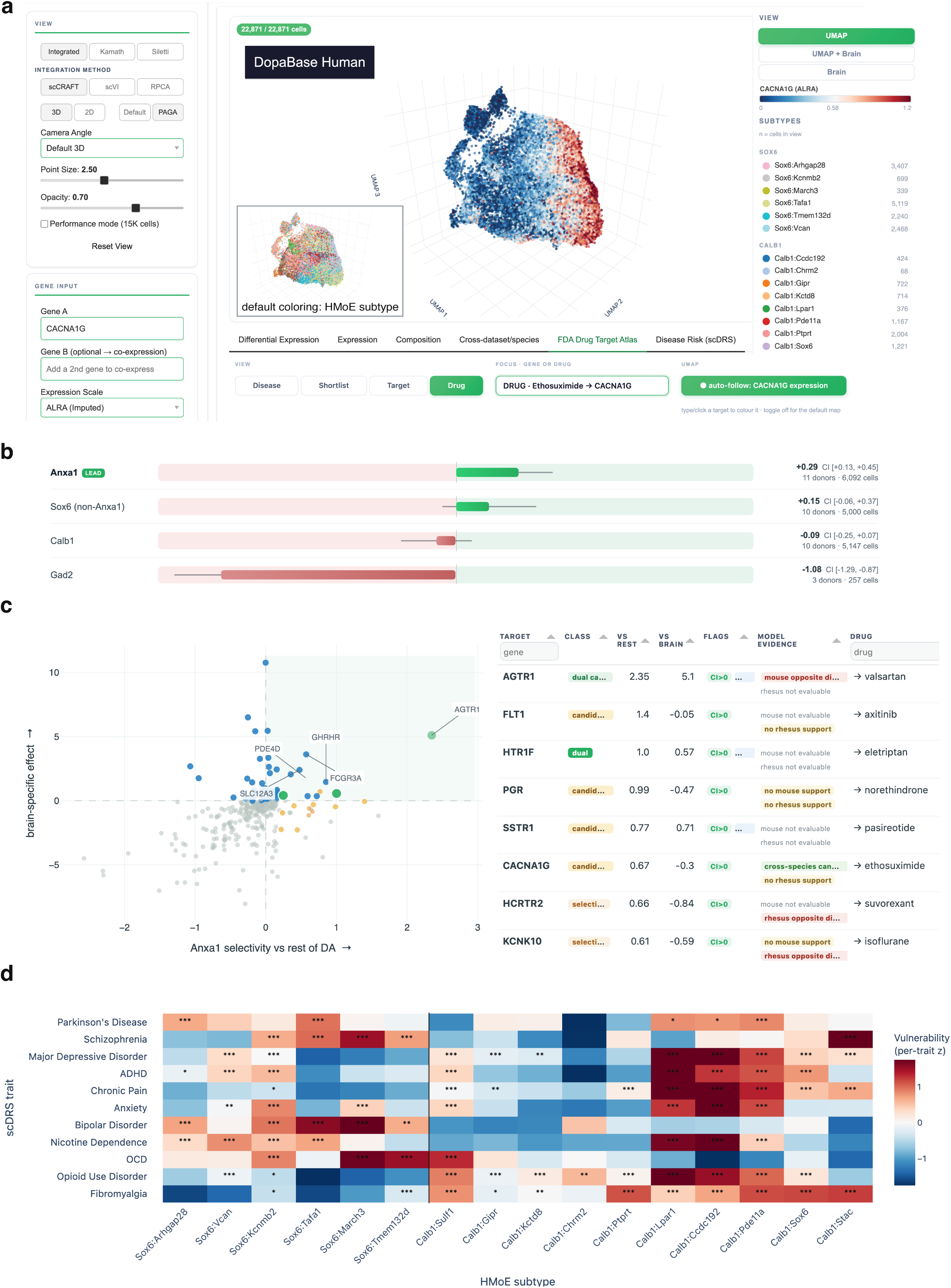
DopaBase Human links dopaminergic subtype identity to disease risk and FDA-regulated pharmacology. **(a)** DopaBase Human view of the integrated Kamath–Siletti atlas containing 22,871 human DA profiles. Selection of ethosuximide identifies its target *CACNA1G* and recolors the three-dimensional UMAP by ALRA-imputed *CACNA1G* expression; the inset shows the default HMoE-subtype coloring. **(b)** PD scDRS enrichment in the Anxa1-associated, Sox6-non-Anxa1, Calb1, and Gad2 populations. Bars show donor-clustered effects relative to the remaining DA population, and whiskers show 95% confidence intervals; contributing donor and cell counts are displayed. **(c)** FDA Drug Target Atlas view for the Anxa1-associated population. The scatterplot places each FDA-regulated target according to its selectivity within DA neurons on the x-axis and its enrichment relative to non-DA populations across the brain on the y-axis. Colors indicate evidence classes; the accompanying table reports effect sizes, detection support, cross-species evidence, and matched FDA-regulated drugs. **(d)** scDRS enrichment across human HMoE terminal subtypes for the eleven GWAS traits displayed in the panel. Values are z-scored within each trait. Asterisks indicate Benjamini–Hochberg-adjusted one-sided Mann–Whitney tests comparing each subtype with the remaining DA cells: *q < 0.05, **q < 0.01, and ***q < 0.001.

Second, for each disease-associated population, we defined an expression-based pharmacological access window by identifying direct molecular targets of FDA-regulated substances according to 1. enrichment over neighboring DA neurons, 2. enrichment over non-DA populations across the brain, and 3. conservation of the corresponding target–population relationship across species. The within-DA comparison identifies targets that distinguish the population of interest from other DA neurons, whereas the whole-brain comparison assesses the specificity of their expression beyond the dopaminergic system. Targets supported by both comparisons were considered dual-selective. Across the atlas, 65 human and 92 mouse target genes were selectively enriched relative to other DA populations in at least one DA population, of which 19 human and 45 mouse targets were also selective relative to non-DA brain populations and therefore met the dual-selectivity criterion. The cross-species axis then determines whether these population–target relationships are conserved between human and mouse, informing how closely the mouse model reproduces the corresponding human molecular selectivity, an important factor for supporting drug testing in mice. Target and drug dossiers show how each target can be perturbed – through agonism, antagonism, or inhibition – alongside blood-brain barrier penetration evidence, disease-genetic evidence, and each compound’s complete direct-target portfolio (**Figure 6c; Supplemental Figure 6a**).

Within the PD-vulnerable Anxa1-associated population, *HTR1F* and *IDH1* are dual-selective in human, whereas *AGTR1* reached the dual-selectivity candidate tier (0.05 ≤ q < 0.10). However, none showed conserved selectivity in mouse: *Htr1f* was not estimable due to dropout, *Idh1* showed the opposite expression relationship, and pooled *Agtr1a*/*Agtr1b* expression was similarly inverted across all mouse donors. This divergence illustrates a central feature of the atlas: human DA-population selectivity, brain-wide specificity, and cross-species conservation are evaluated independently rather than collapsed into a single ranking. By preserving these dimensions separately, the atlas distinguishes targets that are most selective in the human disease-relevant population from those best suited for mechanistic testing across species.

### Population-resolved pharmacology nominates *CACNA1G* as a candidate target in the PD-vulnerable Anxa1-associated population

Calcium-channel blockade is a clinically advanced neuroprotective strategy in PD. Isradipine, the dihydropyridine evaluated in a phase III trial in PD, blocks both CaV1.3 and CaV1.2, encoded by *CACNA1D* and *CACNA1C*, respectively.^97–99^ Neither gene marked the vulnerable Anxa1-associated population. *CACNA1D* was broadly expressed across DA neurons (**Supplemental Figure 6c**), so its expression marks DA identity at large rather than concentrating in the neurons that degenerate first. *CACNA1C* expression was highly localized to a subset of Calb1 family neurons, among the most resilient in PD (**Supplemental Figure 6b**), which places it largely outside the vulnerable population. Instead, *CACNA1G* offered a second route into the same calcium biology, emerging as the strongest cross-species *Anxa1*^+^ candidate: it was near-threshold for enrichment over other human DA neurons and dual-selective in mouse *Anxa1*^+^, with concordant enrichment across species – though it lacked human brain-wide specificity, exhibiting strong expression in the thalamus (**Supplemental Figure 6a**). *CACNA1G* encodes the CaV3.1 T-type calcium channel, which generates low-threshold Ca^2+^ signals selectively in the vulnerable calbindin-negative ventral tier of mouse SNc^100^, making it a pharmacologically tractable target for experimental perturbation. However, existing FDA-regulated compounds engaging CaV3.1 are not *CACNA1G*-selective, illustrating the distinction between target-level selectivity and drug-level specificity captured by the atlas. Ethosuximide engages three targets in the FDA registry. Two had enough detected expression across donors to estimate a within-DA effect, and of these, one was enriched in Anxa1-associated neurons relative to other DA neurons, while neither showed lower expression in Anxa1-associated neurons than in other DA neurons. By comparison, gabapentin engages 34 targets, of which 12 had estimable within-DA effects: four were enriched in Anxa1-associated neurons, and four showed lower expression there than in other DA neurons. Thus, the atlas can identify *CACNA1G* as a selective biological target while simultaneously flagging the polypharmacology of existing compounds that engage it.

### DopaBase Human integrates human DA transcriptomics, disease risk, and pharmacological evidence

To make these analyses accessible and reusable, we developed DopaBase Human, an interactive, open-access resource integrating 22,048 Kamath and 823 Siletti DA profiles into a 22,871-cell human DA atlas. DopaBase Human retains original study annotations, HMoE assignments, and integrated subtype labels in parallel, and supports dataset-specific or integrated 2D/3D UMAP and PAGA views, as well as raw, normalized, or imputed gene-expression visualization (**Figure 6a**).

Within the same environment, users can overlay predicted, integrated, and ground-truth subtype labels on any embedding and query gene expression across them; built-in modules run differential expression, gene-set enrichment, and cross-dataset and cross-species comparisons. Disease Risk views expose the eleven-trait scDRS landscape from family to subtype resolution (**Figure 6d**), while the FDA Drug Target Atlas links implicated populations to dual-axis selectivity, cross-species evidence, and target/drug dossiers (**Figure 6c; Supplemental Figure 6a**). DopaBase Human therefore provides a public analysis framework that moves from a disease-associated DA population to its differentially expressed genes and enriched pathways, and from there to available pharmacology, all accompanied by user-friendly tutorials (**Figure 6c; Supplemental Figure 6e**). By linking disease-associated subtypes directly to FDA-approved pharmacology, DopaBase Human converts subtype vulnerability into experimentally testable hypotheses. Compounds already in human use, such as T-type channel inhibitors acting on CaV3.1, can be prioritized for neuroprotection testing in cellular and animal models.

## Discussion

This work provides seven main advances. 1. A hierarchical mixture-of-experts machine learning model maps the mouse DA subtype organization to human. 2. We identify an Anxa1-associated population that is more severely depleted in Parkinson’s disease than any human subtype described previously. 3. We establish the biological distinctiveness of the Anxa1-associated population across ten complementary analyses spanning prediction confidence, independent classifier convergence, marker expression, depletion relative to its parent populations, size-matched empirical nulls, subtype-agnostic disease-density mapping, PD polygenic risk, cross-dataset and cross-species communication inference, DaT-SPECT, and resting-state fMRI. 4. Ligand-receptor communication affinity analysis, validated against subtype-specific viral tracing in mouse, predicts enriched communication affinity between Anxa1-associated DA and the caudal/posterior putamen. 5. The inferred Anxa-associated putaminal communication affinity field – which resting-state fMRI places in sensorimotor/SCAN cortical territory – is depleted earliest and most severely in 1,127 PD patients scanned by DaT-SPECT, and prodromal risk localizes to this same field within the posterior putamen. 6. We build an FDA Drug Target Atlas using subtype-resolved pharmacology across 11 DA-related diseases, facilitating first-pass screens of any drug targeting the DA system. In PD, a target-first analysis reveals poor alignment between isradipine’s targets and the vulnerable Anxa1-associated population, providing a possible molecular context for its failure to meet endpoints in STEADY-PD III, while prioritizing CACNA1G/CaV3.1 as a cross-species experimental candidate. 7. We release this resource as part of DopaBase Human – a highly detailed, user-friendly portal bridging mouse and human DA gene-expression architecture and connecting disease-associated populations to targets of FDA-approved therapeutics.

Unifying high-resolution subtype identity across mouse and human faces several challenges. Primarily, many genes do not follow the same patterns across species: in mouse, Anxa1 expression defines the subset of DA neurons most vulnerable to PD, but it was not detected at meaningful levels in human when measured by single-nucleus sequencing. Furthermore, age differences across datasets compound the problem – existing mouse DA atlases are composed of cells from young mice, while human DA atlases sequence surviving neurons from postmortem brains where populations may already be depleted. Integration methods designed to remove batch structure can amplify this issue when treating species as the batch – scANVI achieved competitive performance at the family level but assigned only 100 cells to Sox6^Tafa1^, the primary *Anxa1*^+^ subtype in mouse (**Supplemental Figure 2e**). We designed and trained HMoE to address correspondence directly, routing cells through the phylogenetic DA hierarchy and outperforming 13 baseline methods (**Figures 1a and 2b**). HMoE assigned most terminal subtypes with high confidence, including particularly strong predictions for the Anxa1-associated populations, although some subtypes showed broader confidence distributions that likely reflect greater cross-species differences in the organization of the dopaminergic system (**Supplemental Figure 2**). The strong correspondence of the Anxa1-associated populations therefore provided a well-supported basis for asking whether the vulnerability associated with this branch in mouse is also represented in human.

The Anxa1-associated population in humans is depleted in PD well beyond earlier estimates of a most vulnerable subtype (log_2_OR -2.77), surpassing both residual SOX6_AGTR1 (log_2_OR -0.93) and Sox6-family neurons (log_2_OR -1.11; **Figure 3c**). Importantly, HMoE identifies human Anxa1-associated neurons through their broader transcriptional correspondence to the mouse Sox6^Tafa1^ and Sox6^Vcan^ subtypes, rather than an *ANXA1*-expression threshold. Their reduced representation in PD therefore supports selective depletion of this neuronal population rather than merely loss or downregulation of *ANXA1* expression. This heightened differential vulnerability is also reflected in the enrichment of PD risk loci, with GWAS-derived disease scores highest in Anxa1-associated neurons in control donors (**Figure 3a and c**). This is the most severely and selectively affected subset yet identified in human PD, nominating the Anxa1-associated population as a tractable axis for investigating the motor-onset window of PD.

Notably, both the HMoE model and scANVI, an independent reference transfer method, converged on Anxa1-associated DA neurons as a vulnerable subset population that cuts across several annotated human subtypes (**Figure 2e; Supplemental Figure 3g**). Finer clustering of the independently annotated human subtypes might also recover a similarly depleted subpopulation, but it would not by itself establish correspondence to previously characterized mouse populations. Reference-based transfer preserves that correspondence, enabling prior mouse findings^3,16,48^ to inform hypotheses about the human dopamine subtype. One notable result stemming from this transfer is that HMoE assigns ∼35% of DA neurons in the Kamath and Abdulraouf et al. SNc dataset to the Anxa1-associated program (**Figure 2e**), compared with ∼8% of SNc DA neurons expressing *ANXA1* observed directly by RNAscope in age-matched controls^49^, suggesting that marker expression may underestimate the broader transcriptional population in the aged human brain. Establishing this correspondence is therefore critical for translation, because it allows anatomical and behavioral functional features defined experimentally in mice^3,16,48^ to guide hypotheses about analogous human neurons that cannot be directly interrogated in vivo.

Because human dopaminergic projections cannot be traced directly, we inferred preferential striatal targets from ligand-receptor communication affinity, a framework we first validated in mice (**Figure 4c**). This analysis predicts a caudal/posterior putamen region as the most relatively enriched communication affinity target of Anxa1-associated DA neurons (**Figure 4d and e**), converging with prior human diffusion MRI linking lateral SNc to posterior motor putamen and somatomotor cortex.^58,60^ Because those MRI-defined parcels could not determine connection direction or molecular DA identity, our results identify an Anxa1-associated population as a molecular candidate for this macroscale motor nigrostriatal axis – a hypothesis whose disease relevance we then tested against in vivo presynaptic DAT loss.

In DaT-SPECT imaging of 1,127 patients, loss of DAT, which likely reflects both DAT downregulation and axon terminal loss, tracked the Anxa1-associated field specifically, while the Calb1 family was comparatively spared (**Figure 5b–d**).^81^ This territory was already preferentially depleted within the first year of motor symptom onset (**Figure 5b**), consistent with 18F-DOPA PET showing that striatal denervation in early PD begins in the posterior putamen.^101^ Genotype further modified its loss across cross-sectional prodromal and diagnosed cohorts: LRRK2 carriers showed reduced territory DAT before any motor sign, whereas GBA carriers were normal until onset (**Figure 5f**) and then developed an excess deficit that spared the caudate and widened with disease duration, placing GBA-associated vulnerability in the same field (**Figure 5g; Supplemental Figure 5b**). Anxa1-associated field depletion also added prognostic information for prodromal conversion beyond a volume-matched complementary posterior-putamen field, whereas adding the complementary field after the Anxa1-associated field did not (**Figure 5e**).

Paralleling findings in rodent and primate tracing, our affinity maps invert nigral position onto the striatum: Anxa1-associated neurons, the ventral-most tier within Sox6, show their strongest relative affinity for posterior putamen – the primate sensorimotor striatum, homologous to rodent dorsolateral striatum – while dorsally situated Calb1 subtypes favor ventromedial caudate and nucleus accumbens.^1–4,29,51,58,59,102,103^ While ligand-receptor-based inference of communication affinity potential is a proxy for connectivity, and this framework’s concordance with experimental mouse tracing data qualifies it as an inferential first-pass method for use where direct tracing is impossible. Here, we analyze ligand-receptor-derived communication affinity potential, identifying in which striatal regions each subtype’s affinity is relatively greater compared to the rest of DA. These inferred maps should therefore be interpreted as preferential rather than exclusive target fields, consistent with the overlapping connectivity profiles reported for human midbrain subdivisions.^58,104^

Resting-state fMRI further linked this putaminal domain to sensorimotor and somato-cognitive action network cortical territories (**Figure 5h–j**), providing systems-level context for the degeneration of the vulnerable nigrostriatal channel and the early emergence of motor symptoms including bradykinesia in PD.^80^ The emergence of two clinically unique symptom domains, each aligned with a separate DA subtype axis (**Supplemental Figure 5c**), suggests a cellular basis for the heterogeneous presentation of PD across patients.^105–107^ Consistent with a topographic basis for clinical heterogeneity, tremor progression associated more strongly with depletion of the putaminal field inferred to receive Anxa1-associated projections than with depletion of the Calb1-associated field, which spans the ventromedial caudate and nucleus accumbens (**Supplemental Figure 5c**). A recent PPMI DaT-SPECT analysis describes the same relative pattern, linking rest tremor to preservation of caudate dopaminergic terminals with no corresponding difference in the putamen.^108^ Given that fMRI parcellation is population-level, subject-specific differences likely remain, but this approach reflects the overall distribution of functional putamen connectivity.

Dopamine-modulating therapies account for tens of billions of dollars in annual global drug sales^109–111^, yet remain agnostic to molecular DA subtype.^37^ If molecularly distinct nigrostriatal channels contribute to distinct clinical characteristics, then therapeutic targets cannot be evaluated independently of the cellular, anatomical, and species contexts in which they act. For a molecular target, the relevant questions are whether it is enriched in the intended DA population relative to other DA neurons and the rest of the brain, and whether that population–target relationship is conserved in the species used for preclinical testing.^4,37,112–116^ The FDA Drug Target Atlas that we release as part of DopaBase Human addresses these problems by calculating three distinct target-level axes for every direct molecular target of an FDA-regulated substance: selectivity within DA neurons, selectivity relative to the rest of the brain, and conservation across species (**Figure 6c; Supplemental Figure 6a**). At the drug level, the atlas links each compound to its complete direct-target portfolio, allowing its pharmacology to be interpreted in the context of these target-level annotations.

We propose that candidate therapies acting on the DA system should be systematically evaluated across all three dimensions before experimental or clinical prioritization, rather than selected from expression in the intended DA population alone. Such screening can reveal whether a target is enriched in the disease-relevant population, determine whether its expression extends broadly beyond that population, establish whether the same population–target relationship is preserved in the species used for preclinical testing, and identify additional targets engaged by existing compounds that may limit pharmacological specificity.

We identify *CACNA1G* as a cross-species candidate in Anxa1-associated neurons (**Figure 6a and c; Supplemental Figure 6a**), encoding CaV3.1, the predominant T-type channel in SNc DA neurons which mediates low-threshold Ca^2+^ currents.^100,117^ In mice, T-type current density is higher in Calb1-negative ventral-tier than Calb1-positive dorsal-tier SNc neurons.^100^ Their low-threshold depolarizations and dendritic Ca^2+^ signals are reduced by the T-type inhibitor ML218 but not significantly by the L-type blocker nifedipine.^100^ Similar depolarizations occur in *Aldh1a1*^+^ neurons, which include the vulnerable population identified here.^3,100^ Consistent with a role in vulnerability, ML218 reduced rotenone-induced apoptosis in PARK2 patient-derived DA neurons.^118^ These findings support investigating T-type channels alongside the L-type channels targeted by isradipine.

The reasons why isradipine did not demonstrate slowed progression in STEADY-PD III^97^ remain unresolved. Vasodilatory dose limitations may have constrained sustained CaV1.3 inhibition.^99,119^ In addition, chronic genetic loss of CaV1.3 in mouse SNc DA neurons upregulates Cav3.1 mRNA and produces compensatory T-type channel activity^117^, suggesting that T-type channels can take over Ca^2+^-dependent functions when L-type signaling is lost. Our findings raise the possibility that target distribution across DA subtypes may also influence neuroprotective efficacy: neither *CACNA1C* (CaV1.2) nor *CACNA1D* (CaV1.3), the L-type targets of isradipine, was enriched in human Anxa1-associated neurons (**Supplemental Figure 6b and c**). The near-threshold enrichment of *CACNA1G* in human Anxa1-associated neurons, together with concordant mouse enrichment, provides a population-specific rationale for investigating CaV3.1 as an additional target (**Supplemental Figure 6a and d**). The FDA Atlas nonetheless flagged its limited brain-wide specificity (**Supplemental Figure 6a**), consistent with its high thalamic expression and its roles in hypothalamic and prefrontal circuits.^120–122^ CNS-active T-type inhibitors span experimental tools and clinical agents for tremor: ML218 provides selective experimental perturbation, ulixacaltamide met its primary and key secondary endpoints in the Essential3 phase III program and is under FDA review, and zonisamide has been approved as adjunctive PD therapy in Japan since 2009.^123–127^ Together, these data nominate *CACNA1G*/CaV3.1 for direct testing of whether T-type-channel inhibition can preserve the vulnerable Anxa1-associated population and its posterior putaminal field.

Beyond nominating individual candidates, the FDA Atlas lets a user ask, for any one of 11 DA-implicated disorders, which targets enriched in those populations are engaged by existing FDA-regulated compounds, and whether the corresponding subtype–target expression relationship is conserved in the species used for preclinical testing (**Figure 6c and d; Supplemental Figure 6a**) – questions that can expose translational liabilities before substantial preclinical investment. These compounds provide starting points for target interrogation rather than necessarily viable therapeutics. Moreover, the atlas identifies expression-based target selectivity and existing pharmacological relationships; it does not establish target function or therapeutic efficacy, but instead indicates where experimental validation should be directed. Ultimately, via the atlas’s integration with the portal’s suite of interactive analyses, DopaBase Human serves simultaneously as an environment for exploring the major existing human midbrain DA datasets in a single harmonized reference (**Figure 6a**), and as a general engine for mechanism-aware, cross-species hypothesis generation throughout the dopaminergic system.^4,46^

Identifying a highly vulnerable neuronal population, and linking it to the circuit it targets, the genetic risk that accelerates its loss, and the clinical consequences that follow, required experimental access that the human brain does not permit. By resolving this chain computationally – from cross-species subtype identity to inferred targets and in vivo imaging – this work offers a template for studying selective vulnerability of dopaminergic neurons where it ultimately matters but cannot be manipulated: in patients. Extended across diseases and species in DopaBase Human, this foundational platform reframes dopaminergic diseases not as diagnostic categories but as patterns of vulnerability across molecularly defined populations and their circuits. By making these relationships testable in the human brain, this work accelerates the translation of dopaminergic neuroscience into cell-type resolved therapies – and, ultimately, patient care matched to the neurons carrying the largest disease burden.

## Figure Legends

<u>Supplemental Figure 1. Correspondence between source-study annotations and HMoE predictions in an independent mouse atlas.</u> **(a)** Sankey diagram relating the 17 Yaghmaeian Salmani neighborhood annotations to the 18 HMoE-predicted terminal subtypes.^25^ Node height is proportional to cell number, with taller nodes representing subtypes containing a greater number of assigned cells. Ribbons are displayed for mappings containing at least eight cells, and ribbon opacity reflects the fraction of each source neighborhood assigned to the corresponding HMoE subtype. **(b)** Confusion matrix comparing source-study territory annotations with HMoE-predicted families. Fill indicates the percentage of each territory assigned to each family; cells report counts and row percentages, and green outlines identify the dominant predicted family within each territory. Cell-weighted dominant-family concordance was 85.3%.

<u>Supplemental Figure 2. Quality control, independent transfer, uncertainty, marker structure, and comparison with scANVI.</u> **(a)** Characterization of the 1,418 Kamath nuclei assigned to the Gad2 family and excluded from subsequent Sox6/Calb1 analyses. Shown are their position on the Kamath UMAP (i), sequencing-depth and quality-control metrics (ii), dopaminergic and inhibitory marker expression (iii), and the donor-paired DA-program score in retained and excluded nuclei (iv). **(b)** Transfer of the frozen HMoE to 823 midbrain DA neurons from the independent Siletti atlas. Cells are colored by HMoE family (i), HMoE terminal subtype (ii), Siletti subcluster annotation (iii), and an intensity-weighted blend of *SOX6*, *CALB1*, and *GAD2* expression (iv). Profiles identified as DA by the study-independent gate but not assigned to a Siletti DA subcluster are labeled Novel_DA. **(c)** Prediction uncertainty and calibration. Shown are normalized Shannon entropy of the terminal-subtype posterior (i), maximum posterior probability by predicted subtype (ii), and family-level reliability curves for HMoE and scANVI (iii); the diagonal denotes perfect calibration. **(d)** Gene-wise z-scored expression of family- and subtype-associated markers across the Anxa1-associated, Sox6, and Calb1 populations and individual HMoE terminal subtypes. **(e)** scANVI-predicted family (i) and terminal subtype (ii) on the Kamath UMAP. scANVI assigned 100 cells to Sox6^Tafa1^, compared with 4,975 assigned by HMoE, and reassigned much of the corresponding compartment to Sox6^Arhgap^^28^.

<u>Supplemental Figure 3. Population-resolved depletion across PD and Lewy body disease and replication with an independent classifier.</u> **(a–c)** Disease-associated abundance shifts across published Kamath cell types, the HMoE-defined Anxa1-associated population, and its Sox6^Tafa1^ and Sox6^Vcan^ components for PD (a), Lewy body disease (LBD; b), and pooled PD+LBD (c). Analyses included eight control, seven PD, and three LBD donors. Anxa1-associated cells were removed from all other populations before testing. **(d–f)** Corresponding analyses across HMoE-predicted terminal subtypes for PD (d), LBD (e), and pooled PD+LBD (f); the final entry in each family block is the aggregate family estimate, and the Sox6 aggregate excludes the Anxa1-associated subtypes. Points show MASC log2 odds ratios relative to all remaining DA neurons, and whiskers show Wald 95% confidence intervals. Filled points and asterisks indicate Benjamini–Hochberg-adjusted significance: *q < 0.05, **q < 0.01, and ***q < 0.001. **(g)** PD depletion analysis repeated after defining the Anxa1-associated population using scANVI rather than HMoE. Circles denote Kamath reference populations and the square denotes the scANVI-predicted Anxa1-associated population (n=797 PD/control cells assigned to that population). Whiskers show Wald 95% confidence intervals, and a point is starred when its own interval excludes zero. Reference populations exclude the scANVI-predicted Anxa1-associated cells.

<u>Supplemental Figure 4. Communication affinity-derived target fields replicate across independent mouse, macaque, and human datasets.</u> Heatmap values and significance notation are defined as in Figure 4. **(a)** Replication using DA senders from the independent Yaghmaeian Salmani et al. mouse dataset and the same caudoputamen, nucleus accumbens, and olfactory-tubercle receiver regions. **(b)** Communication affinity across caudate, putamen, and nucleus accumbens subregions using macaque midbrain DA senders and basal-ganglia receivers from the Allen HMBA. **(c)** Independent human analysis using Kamath DA senders and Kraft Slide-tags striatal receiver zones. Z1 denotes dorsal striatum enriched regions, Z2–Z3 caudate, Z4 nucleus accumbens, and Z5–Z6 putamen. **(d)** Sensitivity rendering of the human fields using REMAP trained only on the Allen HMBA-BG MERSCOPE reference, rather than the combined MERSCOPE and Slide-tags model used in Figure 4e.

<u>Supplemental Figure 5. Hemisphere-, clinical-, genotype-, and stage-resolved analyses of subtype-weighted DAT loss.</u> MA and LA denote the clinically defined more- and less-affected hemispheres. **(a)** The staging analysis from Figure 5b repeated separately for MA whole striatum (i), LA whole striatum (ii), MA putamen (iii), LA putamen (iv), MA caudate (v), and LA caudate (vi). All matrices use the same color scale and show communication affinity-field-weighted percentage DAT depletion across disease duration, Hoehn and Yahr stage, and MDS-UPDRS part III. **(b)** Hemisphere-resolved genetic analyses. Adjusted SBR estimates for caudate and the Anxa1-associated putaminal field are shown by genotype and hemisphere in (i); estimates were adjusted for age, sex, time since motor onset, and time since diagnosis. Diagnosis-matched contrasts in the Anxa1-associated field are shown in (ii), with negative values indicating lower SBR in the second-named group; whiskers are bootstrap 95% confidence intervals. Spearman correlations between subtype communication affinity fields and voxelwise GBA excess loss are shown in (iii), with spatial block-bootstrap 95% confidence intervals. Corresponding MA and LA GBA excess-loss maps are shown in (iv). **(c)** Clinical associations of subtype-weighted depletion. The difference between Anxa1- and Calb1-field coefficients for longitudinal change in each clinical measure is shown for prodromal participants on the x-axis and de novo PD participants on the y axis (i); colors denote clinical domains. Odds ratios for enrollment iRBD per s.d. increase in field depletion, adjusted for total striatal burden, age, and sex, are shown in (ii). Kaplan–Meier cumulative-conversion curves are shown in (iii) for participants with above-median Anxa1, Sox6, or Calb1 field depletion after residualization against total striatal burden, age, and sex. **(d)** Putaminal SBR differences among LRRK2, GBA, and idiopathic PD across sequential all-visit mixed models. Models progressively include age and sex, time since motor onset, Hoehn and Yahr stage, and levodopa-equivalent daily dose. Red diamonds show the diagnosis-matched primary estimates; negative values indicate lower SBR in the second-named group. **(e)** Age- and sex-adjusted DAT-depletion maps for idiopathic PD (n=937), LRRK2-associated PD (n=124), and GBA-associated PD (n=63), displayed on a common scale. **(f)** DAT-depletion maps across seven intervals of time since motor onset. Sample sizes are shown beside each interval, and all maps use a common color scale.

<u>Supplemental Figure 6. CACNA1G target dossier, calcium-channel expression, and guided analyses in DopaBase Human.</u> **(a)** *CACNA1G* target dossier. The upper section (i) summarizes within-DA and whole-brain effects, dual-selectivity status, FDA-regulated drugs, cross-species estimates, and region-specific expression constraints. The lower section (ii) shows donor-pseudobulk target–disease coupling between *CACNA1G* expression and PD scDRS within the Anxa1-associated population, followed by corresponding coupling estimates across the displayed traits. Coupling is reported as supporting evidence and does not determine target-selectivity class. **(b–d)** ALRA-imputed expression of the L-type calcium-channel genes *CACNA1C*/CaV1.2 (b) and *CACNA1D*/CaV1.3 (c), and the T-type calcium-channel gene *CACNA1G*/CaV3.1 (d), on the Kamath DA-neuron UMAP. Each panel uses an independently scaled color range and should therefore be interpreted spatially rather than compared by absolute color intensity across genes. **(e)** Guided DopaBase Human tutorial for differential-expression analysis, illustrating selection of two cell populations before running the comparison.

## Methods

### Datasets and reference resources

The Gaertner–Oram et al. mouse midbrain snRNA-seq atlas^1^ was the sole HMoE training reference and also supported internal cross-validation and mouse communication-affinity inference. The independent Yaghmaeian Salmani et al. mouse dataset^25^ supported external HMoE validation, paired intact-versus-6-OHDA-lesioned population-depletion analyses, and replication of communication-affinity inference. The Kamath–Abdulraouf et al. human substantia nigra dataset^4^ supported family-level transfer benchmarking, Anxa1-associated population characterization, PD and LBD depletion analyses, and human DA sender profiles. DA neurons from the Siletti whole-brain atlas^46^ provided an independent human transfer dataset. Kamath and Siletti profiles were integrated in DopaBase Human and used for disease-risk and drug-target analyses. No cell from the Yaghmaeian Salmani, Kamath–Abdulraouf, or Siletti datasets entered HMoE feature selection or model fitting at any stage, so every transfer result reported here is out-of-sample.

Whole-mouse-brain MERFISH data from Zhang et al.^128^ supplied mouse striatal receiver profiles. Allen HMBA–basal-ganglia single-cell data^78^ supplied human striatal receivers and macaque DA senders and striatal receivers, and Kraft et al. Slide-tags data^79^ provided an independent human receiver reference. LIANA consensus resources^75^ supplied candidate ligand–receptor pairs. Subtype-specific anterograde tracing maps from Anxa1, Sox6, and Calb1 intersectional driver lines^30^ served as experimental comparators for the mouse target-affinity maps. Allen HMBA–basal-ganglia MERSCOPE^77^ and Kraft Slide-tags^79^ spatial data jointly trained REMAP for spatial rendering. MERSCOPE alone supplied the coordinate reference and the reference-ablation analysis.

PPMI DaT-SPECT images, Xing Imaging Core-Lab binding estimates, and associated genetic and longitudinal clinical records^81^ supported reconstruction validation, DAT-depletion analyses, genotype comparisons, clinical associations, and prodromal phenoconversion modeling. Resting-state fMRI from the NITRC 1000 Functional Connectomes Project^85^ and Human Connectome Project^86^ supported putaminal parcellation and cortical connectivity analyses, which were compared against the somato-cognitive action network reference map of Gordon et al.^67^ Registration and anatomical reference used the symmetric FP-CIT MNI template, the FSL-MNI152 nonlinear 2-mm template, ICBM152-2009 tissue priors, the Oxford-GSK-Imanova structural–anatomical striatal atlas^129^, and HMBA anatomical labels.^77^

GWAS-derived gene sets for eleven DA-related traits, including the Nalls et al. PD GWAS,^74,87–96^ supported scDRS analyses across human, mouse, and macaque DA profiles (**Supplemental Data Table 2**). Non-DA populations of the Siletti whole-brain atlas^46^ and Yao et al. single-cell mouse brain atlas^130^ provided the brain-wide comparators for drug-target selectivity. The FDA Drug Target Atlas integrated GSRS, Drugs@FDA, the Purple Book, openFDA labels, ChEMBL 37, the IUPHAR/BPS Guide to Pharmacology, DrugCentral, Open Targets, HGNC, Ensembl Compara, and B3DB for substance identity, pharmacology, genetic evidence, gene identity, orthology, and brain-penetration annotation. KEGG 2026, Reactome Pathways 2024, WikiPathways 2024 Human, BioCarta 2016, Gene Ontology, and Hallmark collections supported pathway and gene-set enrichment analyses.

### Cross-species dopaminergic subtype prediction

#### Mouse reference dataset

The hierarchical mixture-of-experts model (HMoE) was trained on a published mouse midbrain dopaminergic neuron reference atlas whose annotations were hierarchically organized from three broad molecular families – Sox6, Calb1 and Gad2 – to terminal DA subtypes. These broad molecular families are then further divided into narrower terminal subtype labels. HMoE was trained on this hierarchical taxonomy represented as a rooted tree (**Figure 1a**). The internal nodes within the tree represent the choice of a particular subtype branch for a given DA neuron, whereas the leaf nodes represent terminal DA subtype labels. Lef1 and Gad2^Syndig1^ were excluded because they fall outside the canonical DA hierarchy modeled here, leaving 24,419 neurons assigned to 18 terminal subtypes.^1^

Expression matrices were converted into AnnData objects in Python using scanpy and anndata. Genes from each query dataset were mapped to the HMoE training feature list and ordered identically prior to prediction. Independent mouse datasets were matched based on gene symbols, whereas human genes were mapped to mouse-compatible feature names through ortholog matching. Genes missing from a query dataset were retained in the feature matrix and filled with zeros, allowing each query cell to be evaluated in the same input space as the mouse-trained model.

#### HMoE training and validation

Dopaminergic (DA) subtype identity was predicted with a hierarchical mixture-of-experts (HMoE) classifier implemented in scikit-learn, operating over a fixed, marker-annotated dendrogram of the DA subtypes. At each internal node, an independent expert classifier determined, among the cells reaching that node, which child branch each cell belonged to. Subtype assignment was thereby decomposed into a sequence of biologically structured decisions, from broad DA-family membership down to terminal subtype, rather than a single flat classifier.

The input was raw UMI counts over a fixed training feature set of 21,604 mouse genes. Each internal node was trained independently on only the cells descending from it. Node-specific features were selected among genes detected in at least 0.5% of the node’s cells with a minimum between-branch detection difference of 0.1%, ranked by ANOVA F-statistic (f_classif, sklearn.feature_selection), retaining the top 200 genes per node, selected independently at each node (features not shared across nodes), with training carried to full tree depth. Each node used one of two input representations according to depth within the dendrogram: shallow gates (depth ≤3) used raw UMI counts, whereas deeper gates used analytic Pearson residuals following Lause et al., with a fixed global overdispersion parameter θ=100, clipping at ±√*N*, and non-finite values set to zero.^131,132^ The same transformation was applied during training and inference.

All experts were L2-regularized logistic-regression classifiers (LogisticRegression, sklearn.linear_model) with the lbfgs solver, balanced class weighting, and max_iter = 500, with no additional feature standardization; the inverse regularization strength was C = 1.0 for raw (shallow) gates and C = 0.1 for Pearson-residual (deep) gates. For a query cell, each internal node produced a softmax (probability between 0 and 1) over its children from the corresponding expert logits; these conditional branch probabilities were propagated recursively from the root, multiplied along each root-to-leaf path, summed at the leaves, and renormalized to a distribution over the 18 subtypes. The highest-probability leaf was the predicted subtype, and its probability defined the prediction confidence (**Supplemental Figure 1c**). The finalized model was trained on the complete set of in-model reference cells. Generalization was assessed by 5-fold subtype-stratified cross-validation, in which each fold was trained on four-fifths of the cells and predicted the held-out fifth, so that every cell received exactly one out-of-fold prediction; a single subtype-stratified 90%/10% hold-out provided an additional confirmation. (**Figure 1b; Supplemental Data Table 1**). Performance was summarized at family and terminal-subtype resolution using accuracy, precision, recall, F1, ROC-AUC and precision–recall AUC; all metrics for the 10% held-out set were calculated only from cells excluded during model fitting (**Supplemental Data Table 1**).

#### External validation in the Yaghmaeian Salmani et al. mouse 6-OHDA depletion dataset

To test whether HMoE generalized beyond the training reference, we applied the trained model without retraining to the independent Yaghmaeian Salmani et al. mouse midbrain dataset, comprising 24,094 DAT+ snRNA-seq DA nuclei. Predictions were compared with the source study’s territory and neighborhood annotations and with the established organization of Anxa1-expressing SNc and VTA/dorsal DA populations (**Figure 1c-e; Supplemental Figure 1a and b**). The mouse Anxa1-associated SNc subtype set was defined as cells assigned to the Sox6^Tafa1^ or Sox6^Vcan^ terminal labels. We assessed whether HMoE selectively assigned the SNc *Anxa1*^+^ compartment to these labels while assigning VTA *Anxa1*^+^ DA neurons to their corresponding atlas subtype (**Figure 1f and g**).

### Human transfer

We next applied HMoE to the Kamath et al. human SNc single-nucleus RNA-seq dataset containing 22,048 annotated DA neuron profiles. Human DA neurons were projected into the mouse-trained subtype space after gene-symbol and ortholog harmonization, and HMoE generated terminal subtype probabilities for each cell (**Figure 2a, d-f**). Of 22,048 annotated DA profiles, 1,418 Gad2-like profiles were excluded because they showed reduced dopaminergic identity – and given that the Kamath et al. midbrain dissection is known to be SNc biased – leaving 20,630 cells for Sox6/Calb1 transfer benchmarking and downstream human analyses (**Supplemental Figure 2a**).^4^ To assess transfer to a second human dataset, the frozen HMoE model was applied without retraining to the 823 Siletti midbrain DA neurons – Gad2 predictions were retained because this dataset sampled broader midbrain territories (**Supplemental Figure 2b**). Family-level calibration was assessed by comparing predicted Sox6/Calb1 family identity with observed gene expression accuracy across cells (**Supplemental Figure 2c**).

#### Benchmark comparison

Cross-species transfer performance was evaluated using the independently annotated SOX6/CALB1 family structure in Kamath as an orthogonal benchmark (**Figure 2b and c**). HMoE was compared with 13 baseline approaches: logistic regression, linear support-vector classification, CART decision tree, random forest, naïve Bayes and multilayer perceptron classifiers; nearest-neighbor transfer after PCA or Harmony integration; scVI-nearest-neighbor and scANVI transfer; SingleCellNet; and scGPT representations classified by nearest neighbors or logistic regression. Features (n = 2000)were selected by highest variance in mouse, and classical classifiers and PCA-based methods used counts normalized to 10,000 per cell and log1p-transformed, while scVI and scANVI received raw counts. SingleCellNet performed its package-specific normalization internally, and scGPT used its pretrained-model value-binning procedure. Multinomial logistic regression was fitted using sklearn.linear_model.LogisticRegression with the lbfgs solver, C=1 and max_iter=2000. Log-normalized features were standardized with StandardScaler, fitted only on the mouse reference and subsequently applied to the human query. Sox6 and Calb1 posterior probabilities were obtained using predict_proba. A linear support-vector classifier was fitted using sklearn.svm.LinearSVC with C=1 and max_iter=5000. Features were standardized using the mouse-fitted StandardScaler. Continuous class scores were taken from decision_function and transformed by softmax before Sox6/Calb1 renormalization. The neural-network baseline used two fully connected hidden layers containing 256 and 128 units, respectively, with rectified-linear activation, Adam optimization, and early stopping. Inputs were standardized using the mouse-fitted StandardScaler. Class probabilities were obtained using predict_proba. Random-forest classification used 200 decision trees and the unscaled log-normalized feature matrix. Class probabilities were obtained by averaging tree-level posterior estimates using predict_proba. The CART baseline used an unpruned sklearn.tree.DecisionTreeClassifier with otherwise default package settings and the unscaled log-normalized matrix. Class probabilities were obtained using predict_proba. Gaussian naïve Bayes was fitted using sklearn.naive_bayes.GaussianNB with default package settings and the unscaled log-normalized matrix. Sox6 and Calb1 posterior probabilities were obtained using predict_proba. All unspecified scikit-learn parameters used package defaults. For PCA–k-nearest-neighbor transfer, mouse and human log-normalized matrices were jointly standardized and projected into a shared 50-component principal-component space. A k=15 Euclidean nearest-neighbor classifier was fitted to the mouse coordinates and applied to the human cells using distance weighting. Continuous Sox6 and Calb1 scores were the distance-weighted class fractions returned by predict_proba. This method was transductive because the human cells contributed to the joint scaling and PCA representation. The same jointly standardized 50-component PCA representation was corrected with harmonypy, using species as the batch variable and max_iter_harmony=20. A k=15, distance-weighted Euclidean nearest-neighbor classifier was then fitted on the corrected mouse coordinates and applied to the corrected human coordinates. scVI was trained on raw mouse and human counts under its negative-binomial generative model, with species supplied as the batch variable. Human family identity was transferred from mouse cells in the resulting latent representation using a distance-weighted nearest-neighbor classifier; Sox6 and Calb1 scores were the corresponding continuous class fractions. scANVI was initialized from the scVI model, using mouse family identities as labeled observations and human cells as unlabeled observations. Human class scores were obtained using the package’s soft prediction mode, predict(soft=True), and reduced to Sox6 and Calb1 before scoring. A separate terminal-resolution scANVI model was trained for downstream analyses using the 18 mouse terminal-subtype labels, with mouse cells treated as labeled and human cells as unlabeled (**Supplemental Figure 2e**). As in the HMoE predictions, human cells predicted as Sox6^Tafa1^ or Sox6^Vcan^ were pooled as the scANVI-defined Anxa1-associated population for the terminal-subtype comparison and independent depletion analysis. SingleCellNet was trained on the mouse family labels using its internal library normalization and log1p preprocessing. The package’s continuous SCN_score matrix was used for AUROC calculation. The internal randomized outgroup class was discarded, after which Sox6 and Calb1 scores were renormalized to sum to one. Finally, mouse and human expression profiles were encoded using the same pretrained scGPT model and its native value-binning procedure. The resulting cell embeddings were evaluated through two classifiers. For scGPT–k-nearest neighbors, a distance-weighted nearest-neighbor classifier was fitted on mouse embeddings and used to calculate human Sox6/Calb1 class fractions. For scGPT–logistic regression, a logistic-regression classifier was fitted on the mouse embeddings, and human scores were obtained using predict_proba. HMoE was evaluated by summing terminal-subtype probabilities within the Sox6 and Calb1 branches to generate family-level scores, which were renormalized over those two families before evaluation.

Every method produced a two-column continuous score matrix containing Sox6 and Calb1 scores. Native multiclass outputs were collapsed to these two families, method-specific outgroup scores were removed, and the two retained values were renormalized to sum to one. Accuracy was calculated from the resulting argmax family call. AUROC was calculated from the continuous family score rather than from hard labels. Continuous AUROC inputs were obtained from class posteriors for logistic regression, random forest, CART, Gaussian naïve Bayes, the multilayer perceptron, and scGPT–logistic regression; soft prediction matrices for HMoE, scANVI and SingleCellNet; decision-function margins for the linear support-vector classifier; distance-weighted class fractions for PCA–KNN, Harmony–KNN, scVI–KNN, and scGPT–KNN. Linear-SVM transfer scores represent ranking confidence rather than calibrated probability. Ninety-five-percent confidence intervals were calculated using the Hanley–McNeil closed-form estimator for AUROC and Wilson binomial intervals for accuracy. HMoE was treated as the fixed reference. A baseline was annotated as below HMoE when its upper 95% confidence bound fell below the lower HMoE confidence bound.

### Human Anxa1-associated population characterization

#### Definition of human Anxa1-associated cells

Kamath human SNpc DA neurons were assigned terminal HMoE subtype labels and terminal subtype probabilities as described above. Because human *ANXA1* transcript was not enriched in this population, we use “Anxa1-associated” to denote cross-species alignment to the mouse Anxa1-associated SNc subtype program – Sox6^Tafa1^ and Sox6^Vcan^ – rather than direct human *ANXA1* expression. Of the 20,630 DA neurons retained after classification, 7,256 were assigned to Sox6^Tafa1^ or Sox6^Vcan^ across all diagnostic groups and were defined as Anxa1-associated (**Figure 2e and f**).

#### Marker-based characterization

Enrichment of Anxa1-associated assignments within the published Kamath clusters was summarized using pooled contingency tables. For donor-aware differential-expression analysis, raw counts were summed within each donor for Anxa1-associated and remaining DA cells. Pseudobulk counts were modeled with donor and population terms, log2-fold changes were shrunk using apeglm, and p-values were adjusted by the Benjamini–Hochberg procedure. To identify top detection markers of the Anxa1-associated population, we calculated gene detection differences between Anxa1-associated and remaining DA cells, selecting the top two genes and also reporting shrunk LFCs and adjusted p-values. *PCDH17* and *TPBG* ranked first and second under this criterion; *TAFA1* and *ALDH1A1* were also visualized (**Figure 2f**). Feature plots were generated from ALRA-imputed expression values, with color scales clipped at the 90th percentile. To identify subtype-specific marker genes, we computed rank-based AUROC scores for each gene, equivalent to the normalized Mann–Whitney U statistic, for two contrasts: an individual subtype’s cells vs. remaining within-family DA neurons, and that subtype’s cells vs. all DA neurons (**Supplemental Figure 2d**). Genes were ranked by the minimum AUROC across both contrast scores, prioritizing markers that discriminated the Anxa1-associated population in both comparisons. The subtype heatmap used normalized (10,000 counts per cell), log1p-transformed expression with gene-wise z-scaling.

### PD vulnerability analysis

#### PD-disease relevance scoring

PD-associated transcriptional enrichment was quantified using single-cell disease-relevance scores (scDRS) with gene-level weights derived from the Nalls et al. PD genome-wide association study.^73,74^ Gene-level PD association statistics were obtained by mapping Nalls et al. summary statistics to genes with MAGMA, and the 1,000 genes with the strongest association were retained with their MAGMA z-scores as weights (**Supplemental Data Table 2**). scDRS then scores each cell by the weighted expression of this gene set against 1,000 control gene sets matched for mean expression and variance, so that the score reflects enrichment of GWAS-implicated genes rather than sequencing depth or overall expression level. Scores were calculated for cells from control donors in the Kamath DA dataset and visualized at cell and population resolution (**Figure 3a and c**). For the main population-level display, cell-level scDRS values were modeled as score ∼ population + (1 | donor), a linear mixed model whose donor random intercept matches the structure of the MASC depletion models so that both axes of the figure share one interval convention. Reported effects are contrasts relative to Calb1. Anxa1-associated cells were excluded from the remaining Sox6 and SOX6_AGTR1 comparator populations so that all groups were mutually exclusive. Direct comparisons between Anxa1-associated cells, remaining SOX6_AGTR1 cells and remaining Sox6 cells were performed using donor-level mean scDRS values. For each donor contributing at least ten cells to both populations, the difference in mean score was computed, and these within-donor differences were combined across donors by DerSimonian-Laird random-effects meta-analysis. To evaluate if SOX6_AGTR1 was depleted more than the parent Sox6 family, the same estimator was applied to the entire SOX6_AGTR1 subtype as originally defined, with Anxa1-associated cells retained, against the remainder of the Sox6 family.

#### MELD disease-density visualization

To visualize diagnosis-associated abundance shifts across the continuous transcriptomic manifold without imposing predefined subtype boundaries, Manifold Enhancement of Latent Dimensions (MELD)^133^ was applied to 17,863 DA nuclei from eight control and seven PD donors (LBD cases were excluded). Counts were normalized to 10,000 per cell and log1p-transformed before restriction to control and PD cells; 2,000 highly variable genes were then selected within this subset, scaled with values capped at 10, and reduced to 50 principal components. MELD constructed a k=15 Euclidean neighborhood graph from these components and returned a sample-normalized relative PD likelihood, 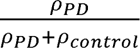. Scores were displayed on the published Kamath UMAP coordinates (**Figure 3d**). MELD was used descriptively, whereas population-depletion inference was performed using the donor-aware MASC models described below.

#### Human and mouse population-depletion analyses

For human depletion analyses, only Kamath control and PD donors were included; LBD was excluded from the primary contrast. In total, eight control and seven PD donors were compared in the primary control-versus-PD analysis (**Figure 3c**). The primary analysis used pairwise-restricted MASC-style binomial generalized linear mixed models. For each focal population, the analysis was restricted to cells belonging to that population or to the Calb1 family, and binary focal-population membership was modeled as 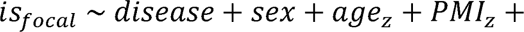 (1 | *donor*). Thus, Calb1 was the literal comparator in each independently fitted model; these were separate binary models rather than a single multinomial model. Sox6 and SOX6_AGTR1 focal populations excluded Anxa1-associated cells, yielding mutually exclusive “remaining Sox6” and “remaining SOX6_AGTR1” groups. Sox6, Calb1, and SOX6_AGTR1 membership was defined from the Kamath annotations, whereas Anxa1-associated membership was defined by HMoE assignment to Sox6^Tafa1^ or Sox6^Vcan^. This analysis was additionally repeated using the scANVI-defined Anxa1-associated population (**Supplemental Figure 3g**). The mouse 6-OHDA depletion was equivalent, except it was modeled between hemispheres, control vs. 6-OHDA depleted, with 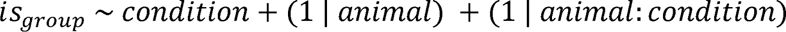 (**Figure 3b**).

Models were fitted using lme4::glmer with a binomial link, the bobyqa optimizer, and maxfun=2*10^5^. Age and postmortem interval were z-scaled, sex was modeled as a factor, and control was the disease reference. Negative disease coefficients therefore indicate lower relative odds of focal-population membership in PD. Effect sizes were reported as log_2_ odds ratios, calculated as the disease coefficient divided by ln(2). Wald 95% confidence intervals were calculated as 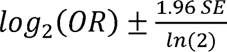, where SE is the standard error of the disease coefficient. Significance of the population estimates was assessed using likelihood-ratio tests comparing the full model with a nested model lacking disease status; the pairwise comparisons were tested at the donor or animal level as described above. Direct comparisons between populations were tested at the experimental unit rather than at the cell, because such a comparison is a within-subject abundance ratio and cell-level tests of it constitute pseudoreplication. For each human donor we computed 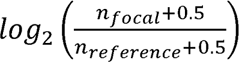 and compared control against PD donors by pooled two-sample t-test, for three comparisons: Anxa1-associated versus remaining SOX6_AGTR1, Anxa1-associated versus remaining Sox6, and remaining SOX6_AGTR1 versus the remainder of Sox6. SOX6_AGTR1 as originally defined, with Anxa1-associated cells retained, was compared against the remainder of the Sox6 family (non-SOX6_AGTR1) by the same test. The mouse analogue used the same ratio computed within each animal and a paired t-test across the six animals. Comparisons against the Calb1 reference are the fitted dots themselves and remained on the glmer. Secondary population-wide analyses used the same human MASC covariates and donor random intercept, but modeled membership in each focal population. Control donors were compared with PD, LBD, and pooled PD + LBD donors (8 control, 7 PD, and 3 LBD) (**Supplemental Figure 3a-f**). Anxa1-associated cells were removed from every other tested population to produce a non-overlapping decomposition, and Benjamini–Hochberg correction was applied across populations within each disease contrast.

To test whether the observed depletion could arise from selecting an arbitrary subset of a vulnerable parent population, 1,000 subsets were sampled without replacement from the Sox6 family and SOX6_AGTR1, matching the number of Anxa1-associated cells within each pool (5,876 of 12,560 Sox6 cells and 4,002 of 7,375 SOX6_AGTR1 cells). Each subset was evaluated using the same Calb1-referenced MASC model. Empirical one-sided p-values were calculated as the fraction of random subsets with depletion at least as extreme as the observed Anxa1-associated subset (**Figure 3e**).

### Cell-to-cell communication-derived striatal target-affinity analysis

#### Ligand-receptor resources and sender-receiver definitions

Dopaminergic–striatal target affinity was inferred using a ligand–receptor tensor framework. Candidate ligand–receptor (LR) interactions were drawn from species-appropriate LIANA resources: the LIANA consensus resource (liana.resource.select_resource(“consensus”)), comprising 4,624 curated LR pairs, for human and macaque analyses, and the LIANA mouse consensus resource for mouse analyses.^75^ Pairs were retained only if both partners were measurable in the analysis data: the ligand had to be detected in the human midbrain DA neuron sender pseudobulk and the receptor in the human striatal receiver pseudobulk. To retain only pairs whose partners vary informatively, both ligand and receptor coefficient-of-variation (CV) filters were applied, retaining pairs whose ligand had CV ≥ 0.3 in log1p pseudobulk expression across the DA sender subtypes and striatal receiver regions.

Five sender–receiver combinations were analyzed: Gaertner and Oram et al. mouse midbrain DA senders with Zhang et al. whole-mouse-brain MERFISH striatal receivers (297 retained LR pairs; **Figure 4a and b**); Yaghmaeian Salmani et al. 6-OHDA mouse DA senders with the same mouse receiver reference (242 pairs; **Supplemental Figure 4a**); Kamath et al. human DA senders with Allen HMBA basal-ganglia striatal receivers (844 pairs; **Figure 4d and e**); Kamath et al. human DA senders with Kraft et al. Slide-tags striatal receivers (837 pairs; **Supplemental Figure 4c**); and Allen HMBA–BG macaque rhesus midbrain DA senders and striatal receivers (840 pairs; **Supplemental Figure 4b**). Species-specific gene symbols were harmonized before construction of each tensor. DA sender populations were analyzed at a resolution of three principal groups: Anxa1-associated cells, the Sox6 family, and the Calb1 family.

#### Communication affinity-tensor construction

The tensor construction and non-negative PARAFAC decomposition follow Tensor-cell2cell, with anatomical receiver bins substituted for the context mode.^75,76^ For each ligand-receptor pair, DA sender subtype and receiver bin, communication affinity scores were computed as:

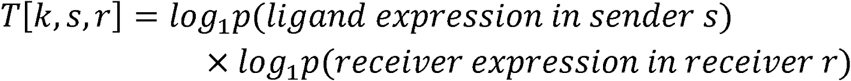

where *k* indexes ligand-receptor pairs, *s* indexes DA sender subtypes, and *r* indexes receiver bins. Expression was summarized from CPM-normalized pseudobulk profiles within each sender or receiver unit. Each ligand–receptor slice was normalized to unit Frobenius norm before tensor decomposition.

#### PARAFAC decomposition and target-affinity reconstruction

Communication affinity tensors were decomposed using TensorLy v0.8.1 non_negative_parafac with rank 20 and n_iter_max = 300. The same decomposition settings were used for the independent mouse, human, and macaque analyses. After decomposition, receiver loadings were L1 soft-thresholded at 0.05. For each sender, the three highest-contributing components were retained to reconstruct an initial target-affinity map. A second within-family PARAFAC pass was then used to resolve target structure among Anxa1-associated, Sox6 family, and Calb1 family sender populations. Reconstructed affinities were converted to per-sender fractions of total affinity across the tested receiver regions or spatial bins. For between-sender comparisons, each fraction was contrasted with the mean fraction of the other two sender groups within the same receiver region such that displayed contrasts represent relative communication affinity enrichment/depletion within each region. Regional enrichment was tested by permuting the three sender-family labels within each receiver bin 5,000 times and recomputing this contrast. P-values were one-sided in the direction of the observed effect and adjusted across receiver regions separately within each sender family using the Benjamini–Hochberg procedure; q<0.05 defined significance. Uncertainty was estimated by bootstrap resampling receiver bins with replacement for 2,000 iterations, with percentile 95% confidence intervals reported descriptively.

#### Validation against mouse tracing maps

Communication affinity-derived target maps were compared with intersectional Cre/Flp dopaminergic-subtype anterograde tracing surfaces from the Anxa1, Sox6 and Calb1 driver lines, matched to the corresponding transcriptomic groups. Tracing surfaces are model-fitted occupancy values on a shared striatal section template, and the tracing map is the inverse-logit fitted eYFP occupancy probability, restricted to template points at which either reporter channel exceeded 0.05 occupancy probability. For each group, both maps were projected onto a shared 16 x 16 dorsoventral-by-mediolateral grid, with the lateral and dorsal directions of the tracing flatmap determined from corticostriatal projections on the same template, where somatomotor cortex marks the dorsolateral extreme. Spatial agreement was quantified as the ROC-AUC with which the continuous communication affinity-derived map discriminates caudoputamen pixels in the upper tertile of tracing intensity, restricted to its largest contiguous region, from the remainder (**Figure 4c**). Rostrocaudal agreement was assessed by placing both modalities on a common bregma axis and comparing mass-weighted profile centers, computed from normalized anteroposterior profiles (section-normalized mean occupancy for tracing, enrichment mass for the communication affinity-derived maps) (**Figure 4c**).

#### Human striatal target-affinity mapping

The canonical human region-level analysis used Kamath DA neurons with HMoE-predicted subtype labels as senders and Allen HMBA–basal-ganglia receiver profiles. The receiver matrix comprised eight regions: body, head and tail of the caudate (CaB, CaH and CaT); core and shell of the nucleus accumbens (NACc and NACs); and caudal, posterior and rostral putamen (PuC, PuPV and PuR) (**Figure 4d**). An independent human analysis paired the same Kamath DA senders with Kraft et al. Slide-tags striatal receiver zones (**Supplemental Figure 4c**). The macaque analysis paired Allen macaque midbrain DA senders with Allen macaque HMBA–basal-ganglia receivers (**Supplemental Figure 4b**).

#### Spatial rendering of human target-affinity fields

Regional human target-affinity estimates were projected into Allen CCF coordinates using REMAP trained on two spatial transcriptomic references: Allen HMBA–basal-ganglia MERSCOPE and Kraft et al. Slide-tags spatial data (**Figure 4e**).^134^ COVET neighborhood representations (k = 30) were learned jointly across both references over 287 shared genes; absolute CCF readout used MERSCOPE alone. The Slide-tags reference therefore shapes the learned neighborhood representation without contributing coordinates. Affinity was scored at each 2 mm voxel from the receiver nuclei REMAP placed there, using the same fitted receiver loadings, and anchored to the eight-region amplitudes so that both readouts sit on one scale. To test how much that contribution matters, we retrained REMAP on the MERSCOPE reference alone and repeated the projection, finding that the rendered fields were robust to dropping the Slide-tags reference (**Supplementary Figure 4d**).

### DaT-SPECT reconstruction and spatial analysis

#### PPMI cohorts and scan selection

DaT-SPECT imaging and clinical data were downloaded from the Parkinson’s Progression Markers Initiative (PPMI) database on 13 July 2026.^81^ The primary baseline analysis included one scan per participant from 230 healthy controls and 1,127 participants with PD for whom a reconstructed volume was successfully registered (**Figure 5a-d**). The PD cohort comprised 937 participants without a recorded LRRK2 or GBA carrier flag, 127 LRRK2 carriers and 66 GBA carriers, including three participants carrying both flags. Double carriers were assigned to the LRRK2 group when mutually exclusive genetic groups were required. SWEDD and prodromal participants were excluded from the primary PD-versus-control analysis.

Quality control required presence in the released Xing Imaging Core-Lab quantitative SBR dataset and successful image reconstruction and registration. Of 1,129 QC-passed PD participants, two images could not be registered, yielding 1,127 baseline PD images. The available longitudinal PD pool contained 2,423 QC-passed scans from 1,129 participants. Baseline analyses used the earliest eligible scan for each participant.

#### DICOM conversion, registration, and preprocessing

Raw HERMES HOSEM Reconstructed_DaTSCAN DICOM series acquired by PPMI were converted to NIfTI using dcm2niix. A scan-specific zero-offset correction was applied by subtracting the first intensity percentile from every voxel, restoring air intensity to approximately zero and correcting integer offsets present in newer acquisitions. Each volume was registered to the symmetric FP-CIT MNI template (symFPCITtemplate_MNI_norm.nii) using a deterministic 12-parameter affine transformation in ANTs with the Mattes mutual-information metric. Registered volumes were resampled to the FSL-MNI152 nonlinear 2-mm template used for the communication affinity-derived maps and smoothed with a 6-mm full-width-at-half-maximum Gaussian kernel.

The cerebral-white-matter reference mask was constructed from the ICBM152-2009 white-matter prior thresholded at 0.9, eroded by two voxels, and excluding a three-voxel dilation of the striatal mask. Reference-normalized voxelwise striatal binding ratios were calculated as

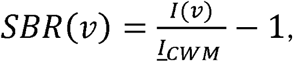

Where *I*(*v*) is the registered uptake at voxel *v* and 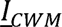 is mean uptake in cerebral white matter.

Regional caudate and putamen measurements used the Oxford-GSK-Imanova structural-anatomical striatal atlas^129^ distributed through NeuroVault image 406338. Posterior, or postcommissural, putamen was defined as the portion of the putamen posterior to the midpoint of the putaminal VOI along the anterior–posterior axis. HMBA–basal-ganglia labels were resampled to the common image space for communication affinity-map weighting and spatial-concordance analyses.

#### Age- and sex-adjusted DAT-depletion maps

Voxelwise expected healthy-control SBR was estimated using ordinary least squares within the control cohort:

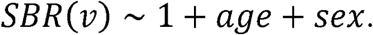

For each PD scan, percentage depletion was calculated relative to the age- and sex-expected healthy-control value:

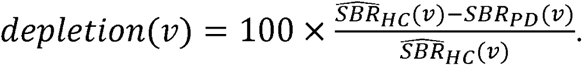

Depletion was calculated only at voxels for which the expected healthy-control SBR exceeded 0.15. Group-level depletion maps were generated by averaging participant-level adjusted maps within the relevant diagnostic or clinical group (**Figure 5a; Supplemental Figure 5e and f**).

#### Validation against Xing Core-Lab SBR values

Reconstructed regional SBR values were compared with the released Xing Imaging Core-Lab cerebral-white-matter-referenced measurements. Validation used one baseline scan from each of 1,350 (1,125 PD, 225 Healthy Control) participants with nonmissing values in both datasets and pooled healthy-control and PD observations. Bilateral whole-VOI means were compared for caudate, putamen, whole striatum, and posterior putamen (defined as the posterior half of the putamen VOI by median-y split, compared against the Core-Lab Postcommissural Putamen measurement) using Pearson correlation, yielding r = 0.81 (caudate), 0.88 (putamen), 0.91 (posterior putamen), and 0.83 (whole striatum).

#### Projection-weighted DAT depletion

Communication affinity-derived target-affinity fields were resampled into the DAT-SPECT image space. Subtype-specific enrichment fields were placed within their corresponding HMBA–basal-ganglia voxels and mirrored across hemispheres to generate bilateral fields. The four tested maps were Anxa1-associated, Sox6-all, Sox6-non-Anxa1, and Calb1.

For participant *i*, projection-weighted depletion was calculated within structure mask *M* as

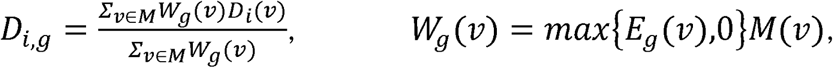

where *D_i_*(*v*) is participant-level percentage depletion and *E_g_*(*v*) is the communication affinity-derived enrichment field for population *g*. Values were calculated separately in the putamen, caudate, and the combined dorsal striatum for all 1,127 baseline PD scans (**Figure 5b and c**).

Population maps were compared within participants using paired Wilcoxon signed-rank tests. Effect size was summarized using matched-pairs rank-biserial correlation, and p-values were Holm–Bonferroni corrected across the six pairwise population comparisons within each structure. Reported percentage-point effects were the median of the within-participant paired differences, not differences between marginal medians. Ninety-five percent confidence intervals for the principal paired differences were estimated using 5,000 nonparametric bootstrap resamples.

#### Voxelwise spatial concordance

Voxelwise concordance was evaluated within a bilateral whole-striatum mask. Projection fields were mirrored from left to right before analysis and compared with the baseline PD-versus-control depletion map using Spearman correlation. The common mask contained 3,211 2-mm striatal voxels. Spatial significance was assessed using BrainSMASH variogram-matched surrogate maps generated with brainsmash.mapgen.base.Base, resample=True, variogram deltas from 0.1 to 0.9, and 1,000 surrogates. Two-sided spatial-null p-values were calculated using BrainSMASH as the prespecified primary spatial null (**Figure 5d**). A companion spatial block bootstrap using 6-mm blocks and 500 resamples was used to estimate confidence intervals and direct differences between map correlations.

#### Disease-duration and clinical-severity analyses

Longitudinal staging analyses included QC-passed PD scans with valid clinical information. A participant could contribute scans from multiple visits. Time since recorded motor onset was divided into [0,0.5), [0.5,1), [1,1.5), [1.5,3), [3,5), [5,8), and [8,40)-year intervals. Clinical severity was examined independently using integer Hoehn and Yahr stages and MDS-UPDRS part III scores grouped in 20-point intervals. To limit domination by densely sampled intervals, sampling was balanced within the finer source bins before adjacent bins were combined into the displayed intervals; exact scan counts are reported in the figure. For each bin, an age- and sex-adjusted group-mean depletion map was generated and summarized within posterior putamen, anterior putamen and caudate VOIs (**Figure 5b; Supplemental Figure 5a and f**). These values are spatial means from group-level maps rather than medians of participant-level regional values.

#### Prodromal phenoconversion and clinical associations

Prodromal analyses used the earliest registered DaT-SPECT scan from 1,665 participants enrolled in the PPMI Prodromal cohort who were free of PD at imaging, had subsequent primary-diagnosis follow-up, and had complete imaging predictors. Participants with PD recorded at or before the baseline scan month were excluded. Phenoconversion was defined as the first primary-diagnosis record of PD after baseline (198 events); participants who did not convert were censored at their last available primary-diagnosis assessment. Posterior putamen was partitioned, using an outcome-independent two-class Otsu threshold, into Anxa1-enriched and complementary territories containing 319 and 369 voxels, respectively. SBR values were sign-reversed and z-standardized so that larger values indicated greater depletion. Phenoconversion was modeled using Cox proportional-hazards regression implemented with statsmodels PHReg and Efron handling of ties. The full model included depletion in both posterior-putamen territories, age, and sex; reduced models omitted either the Anxa1-enriched or complementary territory. Added prognostic information was tested using a one-degree-of-freedom likelihood-ratio statistic, 2(*LL_full_* – *LL_reduced_* (**Figure 5e**). Analyses were performed bilaterally and separately for imaging-defined more- and less-affected hemispheres, with the more-affected side defined as the side with lower baseline posterior-putamen SBR.

Enrollment iRBD (ENRLRBD = 1) was present in 404 of the 1,665 prodromal participants; the remaining 1,261, enrolled for hyposmia or genetic risk, served as the comparison group, and no participants outside the prodromal cohort entered these models. For each whole-striatum projection field, logistic regression modeled iRBD as a function of standardized field depletion, total striatal dopaminergic burden (the pipeline’s whole-striatum mean SBR, sign-reversed and standardized), age, and sex; odds ratios are per standard-deviation increase in field depletion, with Wald 95% confidence intervals (**Supplemental Figure 5c(ii)**). For cumulative-conversion analyses, each field-depletion measure was residualized against total striatal burden, age, and sex, z-standardized, and divided at the median, producing groups of 833 and 832 participants. Kaplan–Meier curves were generated independently for each projection field (**Supplemental Figure 5c(iii)**); participants could therefore belong to the high-depletion group for more than one field, and the 3-year conversion figure is one minus survival at 3 years in the above-median half. Continuous residualized depletion was additionally evaluated using Cox proportional-hazards models adjusted for age and sex, with Efron tie handling; hazard ratios are per standard deviation of residualized depletion with Wald 95% confidence intervals.

Clinical associations were evaluated separately in the prodromal cohort and in de novo PD participants, using 22 clinical assessments grouped for display into motor, other non-motor, neuropsychiatric, and cognitive domains; the groupings were not used in any model. Four were item-level composites of MDS-UPDRS Part III from OFF or unspecified-state visits, with values outside 0 to 4 (including the “unable to rate” code) treated as missing and all component items required: bradykinesia–rigidity (bilateral limb rigidity, finger tapping, hand movements, pronation–supination, toe tapping, leg agility, and neck rigidity; 15 items), rest tremor (five regional amplitudes and constancy), action/postural tremor (four items), and the PIGD score of Stebbins et al. (mean of items 2.12, 2.13, 3.10, 3.11, and 3.12).^135^ Four were MDS-UPDRS or functional totals: the Part IA rater-assessed total, the Part IB patient-questionnaire total excluding pain (sleep, daytime sleepiness, urinary, constipation, lightheadedness, fatigue), the Part II total, and the Schwab–England scale. Pain was analyzed separately as Part IB item 1.9. Thirteen were dedicated instruments: SCOPA-AUT items 4 to 21 and, separately, its swallow/drool items 1 to 3; the Epworth Sleepiness Scale; the RBDSQ (12 symptom items); GDS-15; STAI (state and trait, 40 items); QUIP (13 items); MoCA; Benton JLO; HVLT-R total recall T-score; Letter–Number Sequencing; SDMT; and semantic fluency T-score. Questionnaire sums required at least 75% of items, rounded up (ESS 6 of 8, RBDSQ 9 of 12, QUIP 10 of 13, SCOPA-AUT 14 of 18, GDS-15 8 of 10 forward- and 4 of 5 reverse-scored items, STAI 30 of 40), which affected fewer than 0.5% of records. Cognitive and Schwab–England scores were reversed so that higher values indicate worse function.

For each assessment, the outcome was the change from the baseline score (nearest assessment within 0.5 years of the scan) to the score nearest 3 years after the scan (2-to-5-year window), standardized within cohort. Change was regressed on standardized baseline field depletion, independently reconstructed Xing whole-striatum SBR, the baseline score entered both linearly and as quintile indicators to absorb floor and ceiling effects of bounded items, the exact follow-up interval, age, and sex. Each projection field was fitted in a separate model; the displayed axis is the difference between the Anxa1- and Calb1-field coefficients (**Supplemental Figure 5c(i)**). Sample sizes ranged from 1,039 to 1,171 prodromal and 736 to 772 PD participants across assessments.

#### Genetic PD regional analyses

Genetic analyses used PPMI LRRK2_carrier and GBA_carrier flags defined at baseline. Group membership was fixed across visits, and participants carrying both LRRK2 and GBA variants (3 PD, 16 prodromal) were excluded from genetic comparisons. Baseline models used each participant’s earliest eligible PD scan and the released Xing Core-Lab regional SBR measurements. Idiopathic PD was the reference group. Prodromal genotype comparisons used one scan per participant among healthy controls and prodromal participants with MDS-UPDRS part III ≤ 3. The non-genetic prodromal group included participants enrolled for RBD or hyposmia, and double LRRK2/GBA carriers were assigned to LRRK2. Posterior-putamen SBR was displayed as the raw group mean. Genetic-group comparisons with healthy controls were obtained from ordinary-least-squares models adjusted for age and sex; the non-genetic prodromal group was displayed but was not included in these contrasts.

Posterior-putamen SBR was analyzed using sequential ordinary-least-squares models:

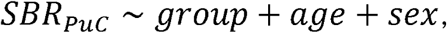

followed by addition of years since motor onset and then Hoehn and Yahr stage. Invalid Hoehn and Yahr values, including negative values and the code 101, were treated as missing. Parallel duration-adjusted models were fitted for caudate, anterior putamen, posterior putamen, posterior putamen and whole putamen. Coefficients are reported in raw SBR units relative to idiopathic PD.

Genotype-specific disease-course trajectories used one baseline scan per prodromal participant and all eligible post-diagnosis visits. Prodromal Anxa1-weighted putamen SBR was summarized across MDS-UPDRS III bins with boundaries at 0, 2, 4, 7, 10, 14, 19, 26, and 34; bins containing fewer than five observations were omitted. Diagnosed scans were grouped in one-year intervals since motor onset, with observations at eight years or later pooled; the prodromal axis was truncated after the last bin in which both carrier groups had at least five participants. Curves show raw means with standard errors of the mean. More- and less-affected hemispheres were assigned from clinical DOMSIDE codes 1 or 2 and fixed across visits; bilateral, uncertain, or missing assignments were excluded. Post-onset decline was summarized as the GBA-minus-idiopathic difference in Anxa1-territory DAT within two years versus beyond four years of onset, tested by patient-level bootstrap (2,000 resamples).

#### Matched genetic analyses

Genetic participants were matched one-to-one to idiopathic-PD participants on exact sex, age within 5 years, and disease duration within 1.5 years. Among eligible controls, the nearest match minimized the sum of the absolute age and duration differences. Matching was performed without replacement. GBA, LRRK2, and combined-genetic analyses were matched independently; the combined set was generated de novo rather than by pooling the separate genotype-specific matches. Posterior-putamen SBR differences were calculated as genetic case minus matched idiopathic control and tested using paired Wilcoxon signed-rank tests. The reported effect was the mean paired difference, with 95% confidence intervals obtained from 5,000 bootstrap resamples of the matched pairs (**Supplemental Figure 5d**).

#### Hemisphere-specific genetic analyses

More- and less-affected hemisphere assignments in genetic analyses were based on clinical DOMSIDE and were therefore independent of the imaging outcome. Baseline caudate and Anxa1-weighted putamen SBR were modeled using ordinary least squares with genotype, age, sex, years since motor onset, and years since diagnosis; adjusted group estimates were predicted at the cohort-average covariate values. Hemisphere-specific diagnosis-matched contrasts used exact matching on sex, age within five years, and time since diagnosis within 0.4 years. Paired differences were tested using Wilcoxon signed-rank tests, with 95% confidence intervals obtained from 2,000 bootstrap resamples (**Supplemental Figure 5b**).

Voxelwise GBA excess loss was calculated as the mean idiopathic-PD minus mean GBA SBR in participants matched on exact sex, age within five years, and disease duration within 1.5 years. Negative voxelwise SBR values were clipped to zero before averaging. Concordance between GBA excess-loss maps and projection fields was quantified using Spearman correlation. Confidence intervals and hemisphere contrasts were evaluated using 500 spatial block-bootstrap resamples with 6-mm blocks; two-sided p-values used the bootstrap standard deviation.

#### Medication-adjusted and longitudinal robustness analyses

For all-visit robustness analyses, whole-putamen SBR was modeled using linear mixed-effects models with participant-specific random intercepts. Four cumulative models included genotype plus age and sex; time since motor onset; Hoehn and Yahr stage; and levodopa-equivalent daily dose (LEDD) (**Supplemental Figure 5d**). LEDD at each scan was reconstructed from the PPMI concomitant-medication interval containing the scan date. Separate longitudinal models among participants with at least two scans modeled posterior-putamen SBR as a function of genotype, disease duration, genotype-by-duration interaction, age and sex, with participant-specific random intercepts and duration slopes. Years since motor onset was centered at 2 years (duration_c_ = duration - 2), so genotype main effects represented group differences in posterior-putamen SBR at 2 years after onset, while genotype-by-duration terms represented differences in annual slope.

### Resting-state fMRI parcellation of the putamen

#### Preprocessing and putaminal functional parcellation

Resting-state fMRI data from 535 healthy participants in the NITRC 1000 Functional Connectomes Project^85^ were used to identify functional subdivisions of the putamen. Images were preprocessed using fMRIPrep^136^, including motion correction, nonlinear registration to MNI standard space, and nuisance regression of six motion parameters, white matter, cerebrospinal fluid, and global signals. For each participant, whole-brain functional connectivity profiles were calculated for individual putaminal voxels. A similarity matrix was constructed from these connectivity profiles and submitted to Louvain community detection^137^. Clustering was performed across network densities of 0.05–0.20 and resolution parameters (γ) of 0.5–2, optimizing community structure by maximizing within-community connectivity and minimizing between-community integration. Individual-level clustering results were combined using consensus clustering, retaining assignments with at least 80% agreement. This procedure identified three bilateral putaminal territories (Put1–Put3).

#### Independent validation of putaminal connectivity networks

The functional connectivity patterns of Put1–Put3 were independently characterized using resting-state fMRI data from 100 healthy participants in the Human Connectome Project^86^. For each participant, the mean BOLD time series was extracted from each putaminal territory and correlated with the time series of all brain voxels. Correlation coefficients were Fisher-z transformed and averaged across participants to generate three whole-brain connectivity maps (N-Put1–N-Put3). These maps characterized the cortical functional networks associated with each putaminal subdivision independently of the dataset used for clustering.

#### Spatial correspondence with the somato-cognitive action network

To determine the relationship between putaminal connectivity networks and the somato-cognitive action network^67^, we quantified the correspondence between the three putaminal connectivity maps and the independently defined SCAN regions. For each participant and putaminal territory, connectivity strength was summarized across voxels belonging to the SCAN reference map. These values were compared across the three putaminal networks to identify the territory exhibiting the strongest functional correspondence with SCAN.

### FDA Drug Target Atlas and DopaBase Human

#### Regulatory-substance, drug and target registry

The FDA Drug Target Atlas was assembled from version-pinned regulatory, pharmacological, genetic and gene-identity resources, including GSRS, Drugs@FDA, the Purple Book, openFDA drug labels, ChEMBL 37, the IUPHAR/BPS Guide to Pharmacology, DrugCentral, Open Targets, HGNC, Ensembl Compara and B3DB. Substance identities were reconciled using GSRS and source-specific identifiers to distinguish regulatory substances, active moieties and marketed products. Ambiguous identity mappings were retained for provenance but excluded from analyses requiring an unambiguous active moiety. The resulting resource contained 3,107 regulatory substances, 2,191 active moieties and 17,722 drug–target relationships.

Drug–target relationships were restricted to direct, primary mechanistic evidence with an unambiguous single-gene target. Indirect pathway relationships, biomarker-only associations, protein complexes without a resolvable primary gene, and nonmechanistic label mentions were excluded from the primary registry. These filters produced a registry of 528 direct, primary, single-gene targets of FDA-regulated substances. Each retained relationship preserved its source evidence, regulatory product, active moiety, mechanism of action, and available potency annotation (**Figure 6c; Supplemental Figure 6a**).

#### Disease-associated DA population mapping

Disease-associated transcriptional programs were mapped across the DA hierarchy using scDRS. Gene sets were obtained from published genome-wide association studies of Parkinson disease, attention-deficit/hyperactivity disorder, anxiety, bipolar disorder, chronic pain, major depressive disorder, nicotine dependence, obsessive-compulsive disorder, opioid use disorder, schizophrenia, and fibromyalgia. For each trait, duplicate-resolved, weighted gene sets of up to 1,000 genes were retained; source studies and final gene lists are reported in Supplemental Data Table 2.

scDRS scores were computed on SCTransform-corrected counts. Variance stabilization was fit separately within Kamath and Siletti, after which the corrected matrices were restricted to their shared genes and scored as a single pooled object, so cells from both studies were ranked on one binning. scDRS preprocessing used 20 mean-expression and 20 variance bins, with an intercept, detected-gene count and donor supplied as covariates; the detected-gene count was taken from raw counts rather than the regularized matrix. Cell scores were calculated using 1,000 matched control-gene sets, mean–variance matching and variance-stabilizing weights, and the normalized score was retained as the reported value. The identical methods were applied in mouse and macaque, substituting the species-specific donor variable. Scores were stored for eleven diseases and exposed at family, original-subtype, HMoE-subtype, and integrated-cluster resolution (**Figure 6b and d**). Group-level summaries were restricted to control donors. DopaBase supported two disease-population definitions. In labeled mode, scDRS enrichment was evaluated in prespecified DA families or subtypes. In unsupervised mode, disease-associated populations were identified from the subtype-level scDRS landscape without prespecifying the focal subtype. Subtypes whose donor-aware mean effect had a confidence interval above zero were included in the disease-associated population.

#### Donor-pseudobulk expression profiles

Target-expression analyses used donor pseudobulks. Raw counts were summed separately within each donor and DA population and converted to log_2_ counts per million. For each focal population, the comparator was the pooled remainder of DA neurons after removal of the focal cells. Only donors containing sufficient cells in both the focal and comparator populations contributed to the paired analysis. Cell-level detection was calculated independently of the pseudobulk effect. The detection difference was defined as the fraction of focal cells expressing the target minus the corresponding fraction among the pooled remaining DA cells. Expression-effect estimates therefore reflected donor-level abundance, whereas the detection criterion required the target to mark a larger fraction of focal cells rather than differing only through expression magnitude in a small subset.

#### Target–disease coupling

Within each species, trait, HMoE population and target, target–disease coupling was evaluated across donor–population pseudobulks. Target expression was calculated as log2 counts per million from raw donor-pseudobulk counts with a prior count of 0.5. Disease burden was summarized as the donor–population mean scDRS normalized score, with the tested target omitted from its own scDRS calculation to avoid direct circularity. Donors contributed with equal weight when at least ten cells were available; analyses required at least three donors, and integrated human analyses additionally required at least two qualifying donors from both Kamath and Siletti. Target expression and scDRS were centered within study × condition × analysis-role strata and independently z-standardized. Their association was estimated as a standardized linear coefficient, with degrees of freedom equal to the number of donors minus the number of strata minus one. Two-sided t-based 95% confidence intervals and p-values were calculated, and p-values were Benjamini–Hochberg-adjusted across FDA targets separately within each species × trait × population × analysis partition × scoring method. Coupling estimates were presented as independent supporting evidence and did not contribute to target-selectivity classification or ranking (**Supplemental Figure 6a(ii)**).

#### Within-DA target selectivity

Within-DA selectivity was defined as the donor-paired mean log2-CPM difference between the focal population and the pooled remainder of DA neurons. Enrichment was tested using a one-sided paired t-test on donor-level log2-CPM differences, with confidence intervals calculated from the paired donor effects. Benjamini–Hochberg multiple-testing correction was performed within the estimable FDA-target panel for each population.

A target was classified as selective when: *q*_FDA_ < 0.05, CI_lower_ > 0, and a detection difference ≥ 10%. A selectivity candidate met the confidence-interval and detection criteria but had an FDA-target-family q-value between 0.05 and 0.10. Candidates were therefore near-misses on the prespecified FDR criterion rather than nominal signals lacking effect or specificity support (**Figure 6c; Supplemental Figure 6a**). Targets with a positive, confidence-supported expression effect but broad detection in the focal population were classified separately as graded when they were detected in more than 80% of focal cells and had a detection difference below 0.10. This category captured targets following a continuous lineage or vulnerability gradient rather than marking a discrete population. Populations represented by fewer than four common donors were designated descriptive only. No confidence-gated selective, candidate, or graded classification was assigned to these populations. The same rules were applied in human, mouse, and macaque. Calibration of the FDA-target-family procedure was evaluated using donor sign-flip permutations.

#### Non-DA whole-brain selectivity

Brain-wide analysis tested whether within-DA target enrichment remained selective beyond the dopaminergic system. For the human analysis, focal DA expression incorporated all 11 DA control donors represented in the integrated Kamath–Siletti dataset. The reference consisted of whole-brain non-DA populations after removal of 2,026 putative DA cells identified by the union of four independent DA definitions. The same donor-aware estimator and target-family correction were applied to human and mouse brain-wide analyses. Brain-wide effects were expressed as mean log2-CPM differences between the focal DA population and the non-DA whole-brain reference. Region-specific contrasts were also retained so users could determine which brain region imposed the strongest constraint on selectivity or required specified off-population regions to lack competing enrichment.

A target was classified as dual-selective when it passed the within-DA selective gate and also showed a positive, FDR-significant brain-wide effect. A dual-selectivity candidate met the within-DA selectivity-candidate criteria but was significantly enriched relative to the whole-brain reference. Thus, dual selectivity required support on two distinct axes: discrimination from neighboring DA populations and restriction relative to non-DA populations across the brain (**Figure 6c; Supplemental Figure 6a**).

#### Cross-species classification

Following ortholog harmonization, to avoid systematically excluding clinically relevant targets with one-to-many mouse orthology, for human targets with multiple mouse orthologs, raw mouse counts were summed across all mapped paralogs before pseudobulk construction. The final human–mouse conservation universe contained 511 evaluated target loci. Cross-species evidence was summarized using three manuscript-facing labels: a) conserved, for target–population relationships with concordant, supported enrichment in human and mouse/macaque; b) conserved candidate, for directionally concordant relationships with strong evidence in one species and compatible but incomplete support in the other; and c) unresolved or divergent, for relationships with opposing directions, inadequate coverage or insufficient evidence for a concordant classification. A separate magnitude-heterogeneous flag identified directionally concordant relationships whose effect sizes differed substantially between species. “Selectivity candidate” refers to within-species target evidence and is distinct from the cross-species label “conserved candidate” (**Figure 6c; Supplemental Figure 6a(i)**).

#### Three-species target prioritization

Across all focal population–target combinations with estimable human, mouse, and rhesus effects, cross-species evidence was combined using fixed-effect inverse-variance meta-analysis. Positive enrichment was prespecified. Targets were retained when effects were concordant in direction across all three species, the one-sided meta-analysis passed Benjamini–Hochberg adjusted q<0.05, and Cochran’s Q showed no significant between-species heterogeneity after correction. Corrections were performed separately within each focal population across the estimable FDA-target panel. Retained targets were ranked by the minimum species-specific Cohen’s h, a standardized measure of focal-versus-comparator detection enrichment, thereby prioritizing targets with consistently selective detection across all three species.

#### Drug mechanism and target-portfolio annotation

Mechanism-of-action annotations were harmonized across the pharmacological and regulatory sources into three source-reported directions (activation, inhibition, ambiguous), then resolved per drug–target pair into a single consensus label: agonist, inhibitor, mixed (conflicting activation and inhibition), modulator (direction reported only at broad target scope) or binder (no directional action available). Source-reported potency values were retained with their assay type and units; potency values from different assay systems were not treated as directly interchangeable. Approved indications, regulatory status and available blood–brain barrier or CNS-penetration evidence were attached at the active-moiety level. Disease-genetic evidence was summarized independently from expression selectivity using fine-mapped credible sets, QTL colocalization and Open Targets genetic evidence. This allowed the Atlas to distinguish a target that selectively marked a disease-associated population from a target independently implicated as a disease-risk gene.

Target selectivity was defined for each gene within a focal DA population and did not imply that every drug engaging that target was itself population-selective. Drug dossiers therefore listed each compound’s complete set of direct registry targets and, for the selected DA population, summarized how many measurable targets were selectively enriched, nonselective, or negatively enriched. These descriptive portfolio counts indicated whether the compound’s additional targets supported or weakened the population specificity suggested by its best-ranked target; they did not create a separate statistical tier. For shortlist navigation, DopaBase ordered candidates by measured selectivity alone. Candidates were sorted first by selectivity tier (dual-selective, selective, dual candidate, borderline) and then by the user-selected ranking axis: within-DA enrichment percentile, whole-brain percentile, or a joint rank defined as the minimum of the two direction-aware percentiles, so that a weak axis cannot be masked by a strong one. Disease vulnerability, human genetics, cross-species conservation, mechanism direction, clinical development stage, and CNS penetration were identified as independent filters and presented as evidence, but none were included in the ordering. Shortlist membership was determined solely by focal-versus-pooled-rest-of-DA enrichment under the shared FDA-family FDR rule.

### DopaBase Human construction and interactive analyses

#### Dataset assembly and harmonization

DopaBase Human integrated 22,048 DA profiles from the Kamath substantia nigra dataset with 823 DA neurons selected from the Siletti whole-human-brain atlas, yielding 22,871 cells across 25,037 shared genes (**Figure 6a**). Siletti cells were obtained from the CZ CELLxGENE Census release and restricted to neuronal profiles from substantia nigra, periaqueductal gray, and red-nucleus dissections. Candidate DA cells were identified using a two-component Gaussian-mixture model applied to a machinery score calculated from *TH*, *DDC*, *SLC6A3* and *SLC18A2*. Cells were additionally required to express at least two of these four genes; serotonergic and noradrenergic profiles were removed using *TPH2*- and *DBH*-based thresholds.

Siletti source-subtype annotations were recovered by mapping the selected cells to the author-annotated Siletti DA reference. After normalization and principal-component analysis, each query cell was assigned the majority subtype among its 15 cosine-nearest reference neighbors when the nearest-neighbor correlation was at least 0.95; otherwise, it was labeled Novel_DA. Before integration, gene symbols were converted to uppercase, duplicate symbols were summed, and the two datasets were restricted to shared genes. Raw integer UMI counts were retained. Study, donor, disease status, anatomical region, and original subtype metadata were harmonized before concatenation.

#### Integration and annotation

The primary integrated representation was generated using scCRAFT, with study of origin used as the batch variable (**Figure 6a**). Genes detected in fewer than five cells were excluded, counts were normalized and log-transformed for integration, and 2,000 highly variable genes were selected after excluding mitochondrial and ribosomal genes. The scCRAFT model was trained for 150 epochs, and the learned representation was reduced to 50 dimensions for downstream neighborhood construction. scVI and Seurat v5 reciprocal-PCA integrations were generated as alternative views. scVI used study as the batch variable and a 30-dimensional latent representation. Reciprocal-PCA integration used 3,000 integration features, Kamath as the reference dataset, dimensions 1–30, and k.anchor=5. A 15-nearest-neighbor graph was constructed in the 50-dimensional scCRAFT representation. Initial Leiden over-clustering was performed at resolution 1.2, after which clusters were iteratively merged until every remaining pair differed by at least ten genes with an absolute log2 fold change greater than 1 and Benjamini–Hochberg FDR below 0.05, yielding 17 integrated clusters.

Cluster names were assigned using count splitting to separate clustering from marker selection. Raw counts were divided by binomial thinning into training and held-out matrices, and differential expression on the held-out counts was then used to name each cluster by its DA family and highest-ranked non-housekeeping marker. Original Kamath and Siletti subtype labels, HMoE subtype and family assignments, and integrated-cluster annotations were retained as parallel metadata fields rather than reconciled by voting. The manuscript refers to the named integrated Leiden clusters as integrated subtype annotations. Broad family labels were assigned hierarchically. A three-component Gaussian-mixture model identified *GAD2*-high cells; the remaining cells were assigned to Sox6 or Calb1 according to relative standardized *SOX6* and *CALB1* expression.

#### UMAP, PAGA and anatomical visualization

Two- and three-dimensional UMAP embeddings were precomputed for the scCRAFT, scVI and reciprocal-PCA representations using 15-nearest-neighbor graphs (**Figure 6a**). PAGA connectivity was calculated over the corresponding integrated clusters. Because Scanpy’s standard PAGA initialization is two-dimensional, we implemented a custom three-dimensional PAGA initialization. The PAGA cluster graph was laid out in three dimensions by classical multidimensional scaling of its shortest-path distances, each cell was initialized at its cluster’s node position with small isotropic jitter, and a three-component UMAP was then optimized from these coordinates. This preserved the PAGA-derived global arrangement while permitting local UMAP organization in three dimensions. Dataset-specific views were also retained. The Kamath view included the published two-dimensional coordinates and a reproduced LIGER latent representation for additional embeddings. The Siletti-specific view used principal-component analysis followed by BBKNN with donor as the batch variable.

#### Expression visualization and differential expression

Gene expression could be viewed as raw counts, total-count-normalized and log1p-transformed expression, or ALRA-imputed expression (**Figure 6a**). ALRA factors were precomputed, whereas raw and normalized gene vectors were calculated on demand. DopaBase Human provides separate cell-level and donor-pseudobulk differential-expression modes. Cell-level analysis uses Scanpy Wilcoxon rank-sum tests after total-count normalization and log transformation. Genes are retained when detected in at least 10% of either group with a mean raw count greater than 0.1 in either group. Because cells rather than donors were the observational units, cell-level results were intended for exploratory analysis. For donor-pseudobulk analysis, raw counts were summed separately within each donor and selected group. A donor contributed when at least ten cells were available for the relevant group, and each comparison required at least two qualifying pseudobulks per side. When at least two donors contributed to both groups, PyDESeq2 models included donor and group terms and were restricted to the shared donors; otherwise, an unpaired group-only design was used. Wald statistics and log2 fold changes were reported, and p-values were adjusted using the Benjamini–Hochberg procedure. The guided implementation of this comparison is illustrated in **Supplemental Figure 6d**.

#### Gene-set enrichment and composition analysis

Preranked gene-set enrichment analysis used GSEApy with genes ordered by the differential-expression test statistic: the DESeq2 Wald statistic for donor-pseudobulk analysis and the Wilcoxon z statistic for cell-level analysis. Analyses used 1,000 permutations, and normalized enrichment scores and FDR q values were reported. Over-representation analysis was performed separately for positively and negatively differentially expressed genes meeting Benjamini–Hochberg-adjusted p≤0.05. The background consisted of all genes tested in the differential-expression comparison. Gene sets were tested using one-sided Fisher exact tests, with separate Benjamini–Hochberg correction for the positive and negative directions. For both preranked and over-representation analyses, each gene set was first intersected with the genes tested in that comparison and was evaluated only if 15 to 500 of its genes remained. The default human canonical-pathway collection combined KEGG 2026, Reactome Pathways 2024, WikiPathways 2024 Human, and BioCarta 2016; Gene Ontology and Hallmark collections were also selectable. The composition module displayed the fraction of each donor’s selected cells (normalized within donor) assigned to each population label.

#### Disease-risk and cross-dataset analyses

scDRS scores were precomputed separately within Kamath and Siletti because the scoring expression matrices were not corrected across sequencing platforms. Expression was normalized with SCTransform, fit separately within each study. scDRS preprocessing used 20 mean-expression and 20 variance bins with covariates of an intercept, detected-gene count, and donor. Detected-gene count was taken from the raw matrix rather than the SCT matrix, which is regularized and would understate sequencing depth. Cell scores were calculated using 1,000 matched control-gene sets, mean–variance matching, and variance-stabilizing weights. Scores were stored for eleven diseases and exposed at family, original-subtype, HMoE-subtype, and integrated-cluster resolution. The application provided per-cell distributions, group-level summaries, heatmaps, and stratified disease-risk views (**Figure 6b and d**). Heatmap inference compared each population with the remaining cells using one-sided Mann–Whitney tests followed by Benjamini–Hochberg correction within each trait.

Kamath–Siletti annotation correspondence was displayed through direct cross-tabulation when both labels were available on the same cells. Otherwise, expected correspondence was estimated through the shared integrated-cluster annotation; these mediated values were explicitly displayed as expected co-abundance rather than observed cell counts. Human–mouse subtype correspondence was computed by correlating homolog expression profiles between species: for each subtype in each species, mean log-normalized expression was calculated across one-to-one orthologs, and cross-species pairs were scored by Spearman correlation, with each subtype assigned to its top-correlated counterpart.

#### Application implementation and release validation

DopaBase Human was implemented in Python using Panel and Bokeh and deployed as a Docker application. The integrated AnnData object, embeddings, ALRA factors, anatomical meshes, scDRS values, and FDA Atlas tables were precomputed. User-defined differential-expression and enrichment analyses are calculated on demand.

### Statistical analysis and reproducibility

No statistical method was used to predetermine sample size. Analyses were performed in Python v.3.10.19 and R v.4.5.0 using packages and versions as follows. Python: numpy 2.2.6, pandas 2.3.3, scipy 1.15.3, scanpy 1.11.5, anndata 0.11.4, scikit-learn 1.7.2, torch 2.5.1, tensorly 0.8.1, statsmodels 0.14.6, scdrs 1.0.2, and matplotlib 3.10.8. R: Seurat 5.4.0, SeuratObject via Matrix 1.7-3, sctransform 0.4.2, lme4 1.1-37, metafor 4.8-0, and ggplot2 4.0.2. Exact sample sizes and statistical tests are reported in the corresponding Methods sections and figure legends.

## Availability statement

Code is available at https://github.com/AwatramaniLab/schonfeld_etal_anxa1_2026 and archived at https://zenodo.org/records/22863841. The GWAS-derived gene lists used for the scDRS analyses (also provided in Supplementary Data Table 2), together with all code and models generated in this study, are deposited at Zenodo and listed with their persistent identifiers in the Key Resources Table. The HMoE model is available to annotate user supplied sequencing data from dopaminergic neurons at https://huggingface.co/spaces/awatramani-lab/hmoe-da, and weights are open-sourced at https://huggingface.co/awatramani-lab/hmoe-da. DopaBase Human will be made publicly available at time of publication. No new data/protocols were generated in this study.

## Acknowledgments

This research was funded by Aligning Science Across Parkinson’s ASAP-025188 (Team Dombeck) and ASAP-027450 (Team Awatramani) through the Michael J. Fox Foundation for Parkinson’s Research (MJFF). R.A. was also supported by a grant from the National Institutes of Health (2R01NS119690-06). For the purpose of open access, the author has applied a CC BY public copyright license to all Author Accepted Manuscripts arising from this submission.

## Contributions

ES performed all transcriptomic and DaTscan analyses depicted, developed the HMoE model, and wrote the manuscript. RA supervised the study. MB and AA provided the fMRI analyses. JS, DD, BY, TP, JP, MM, and TH provided critical feedback on transcriptomic analyses, human *Anxa1* expression, calcium channels in neurodegeneration, selective vulnerability, PPMI patient cohorts and interpretation. All authors helped revise the manuscript.

