## Supplementary figures and images for "Identification of a human Anxa1-associated dopaminergic population linked to Parkinson’s disease vulnerability and sensorimotor putaminal networks"

### Supplemental Figure 1

#### Supplemental Figure 1

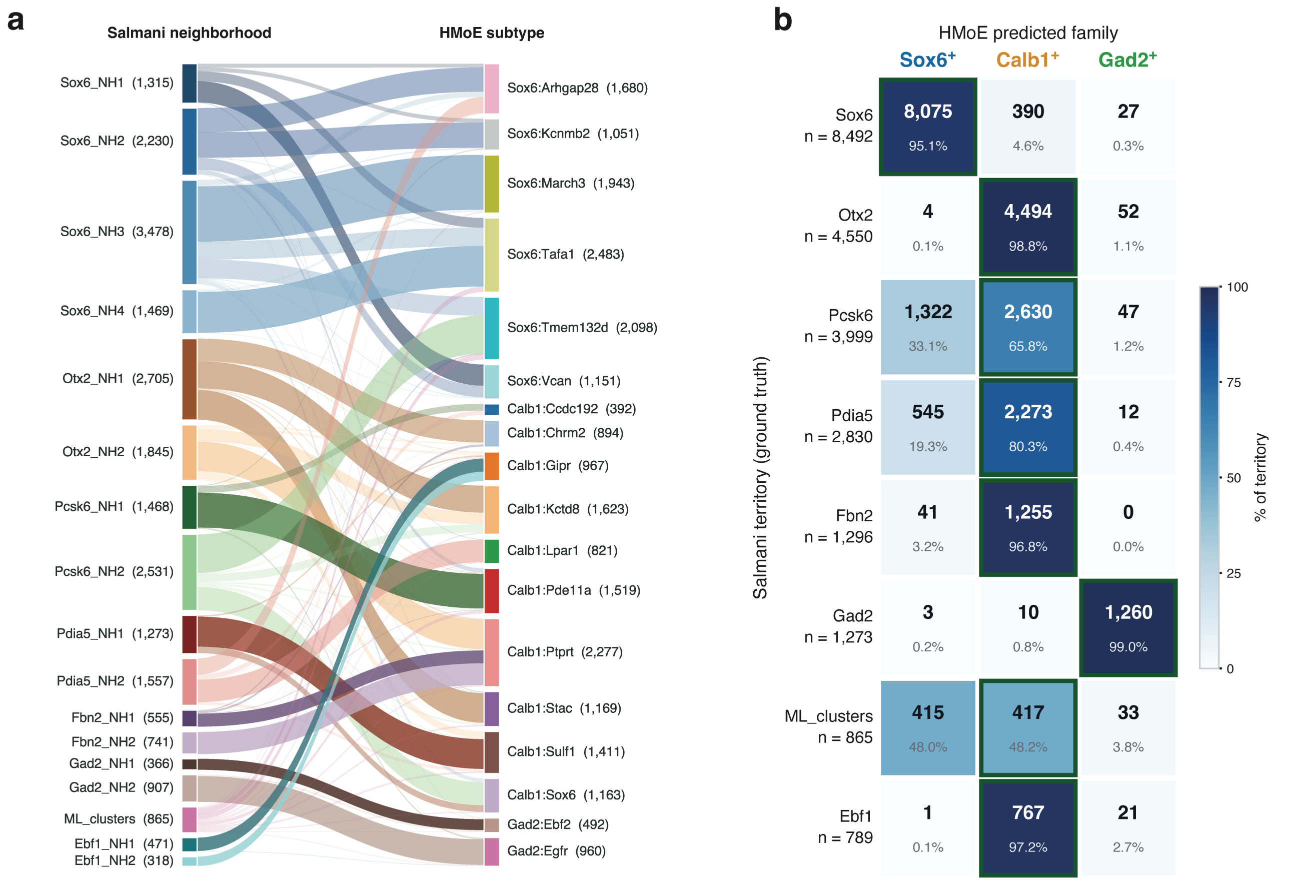

### Supplemental Figure 3

#### Supplemental Figure 3

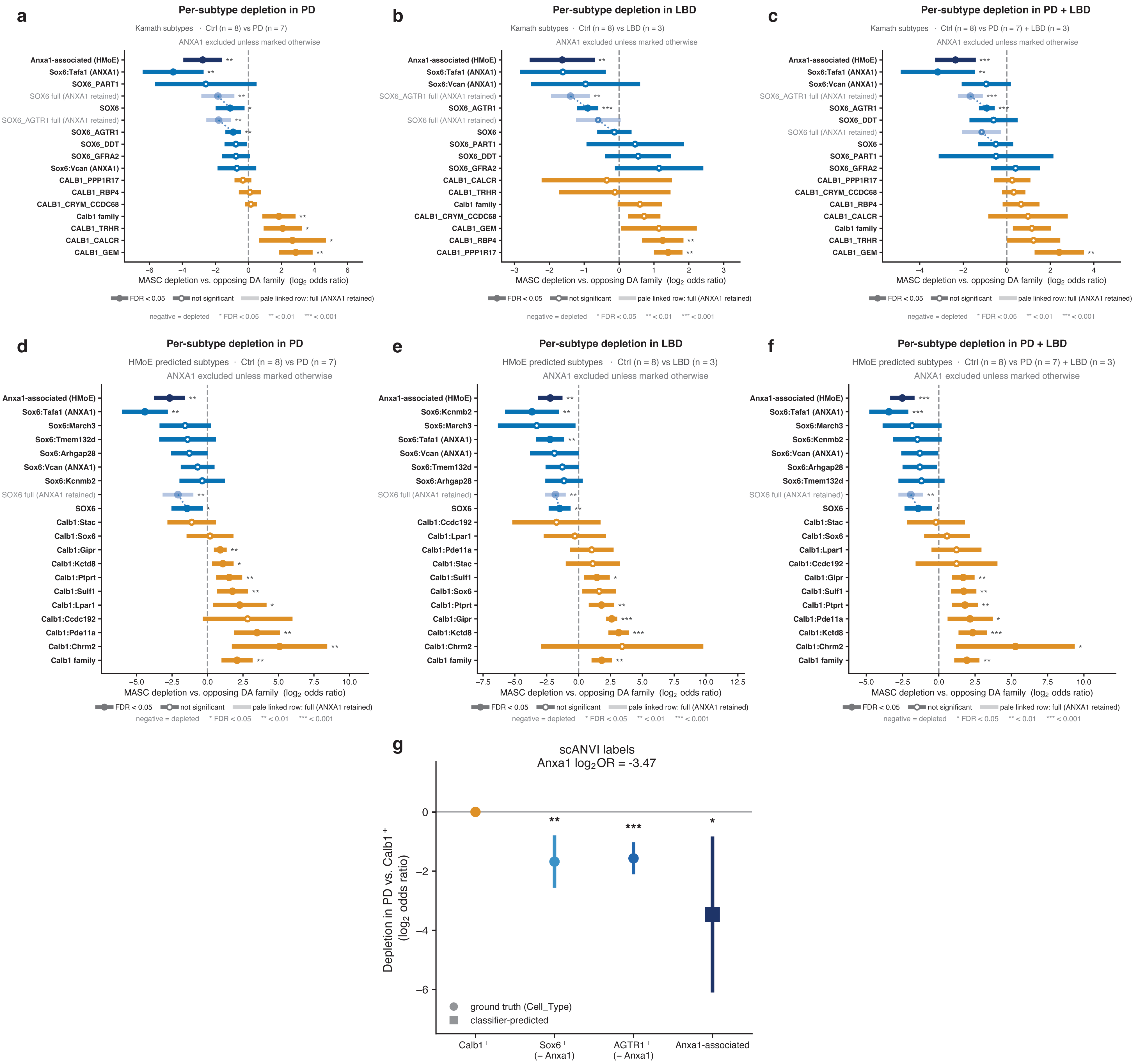

### Supplemental Figure 4

#### Supplemental Figure 4

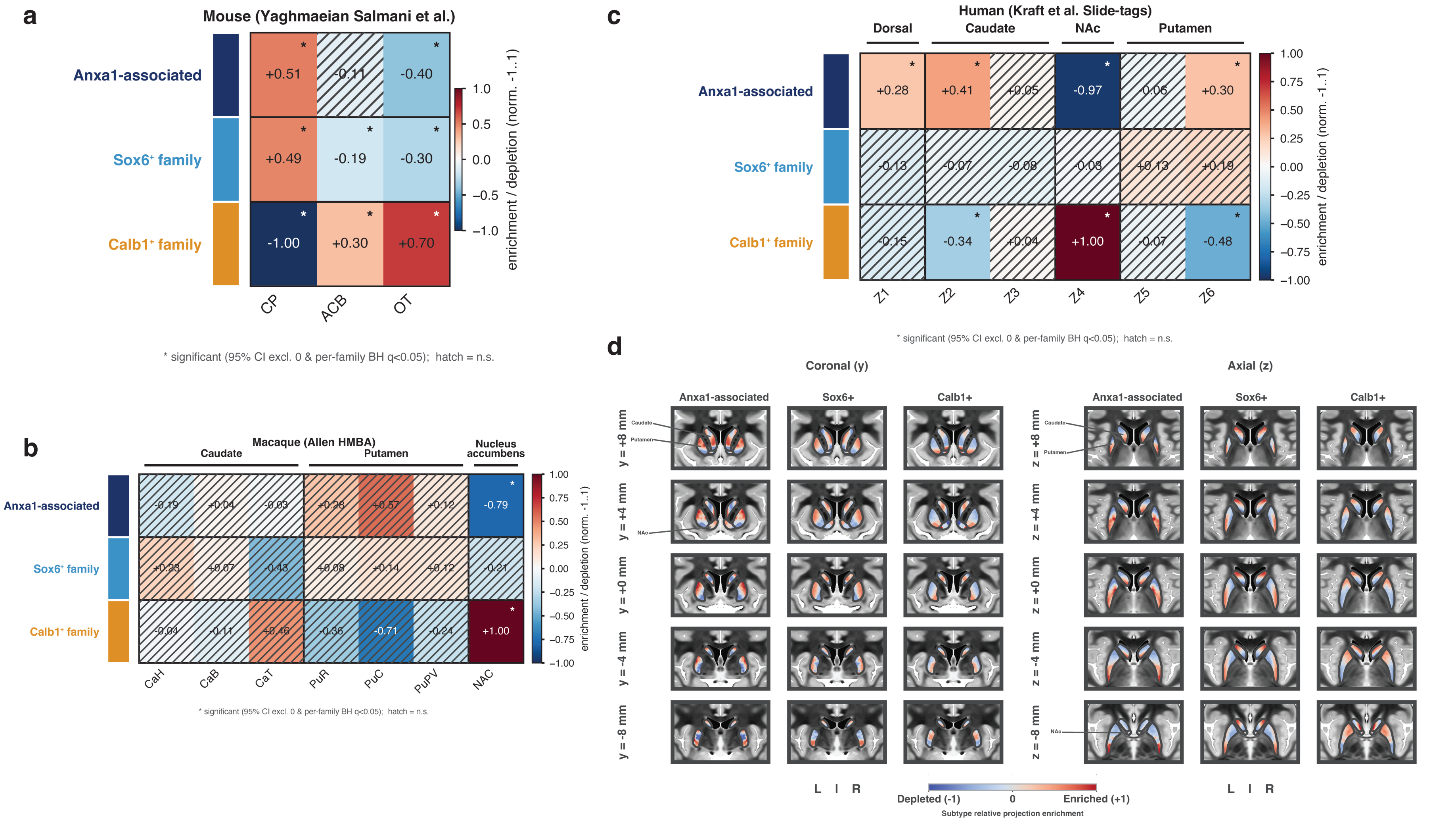

### Supplemental Figure 5

#### Supplemental Figure 5

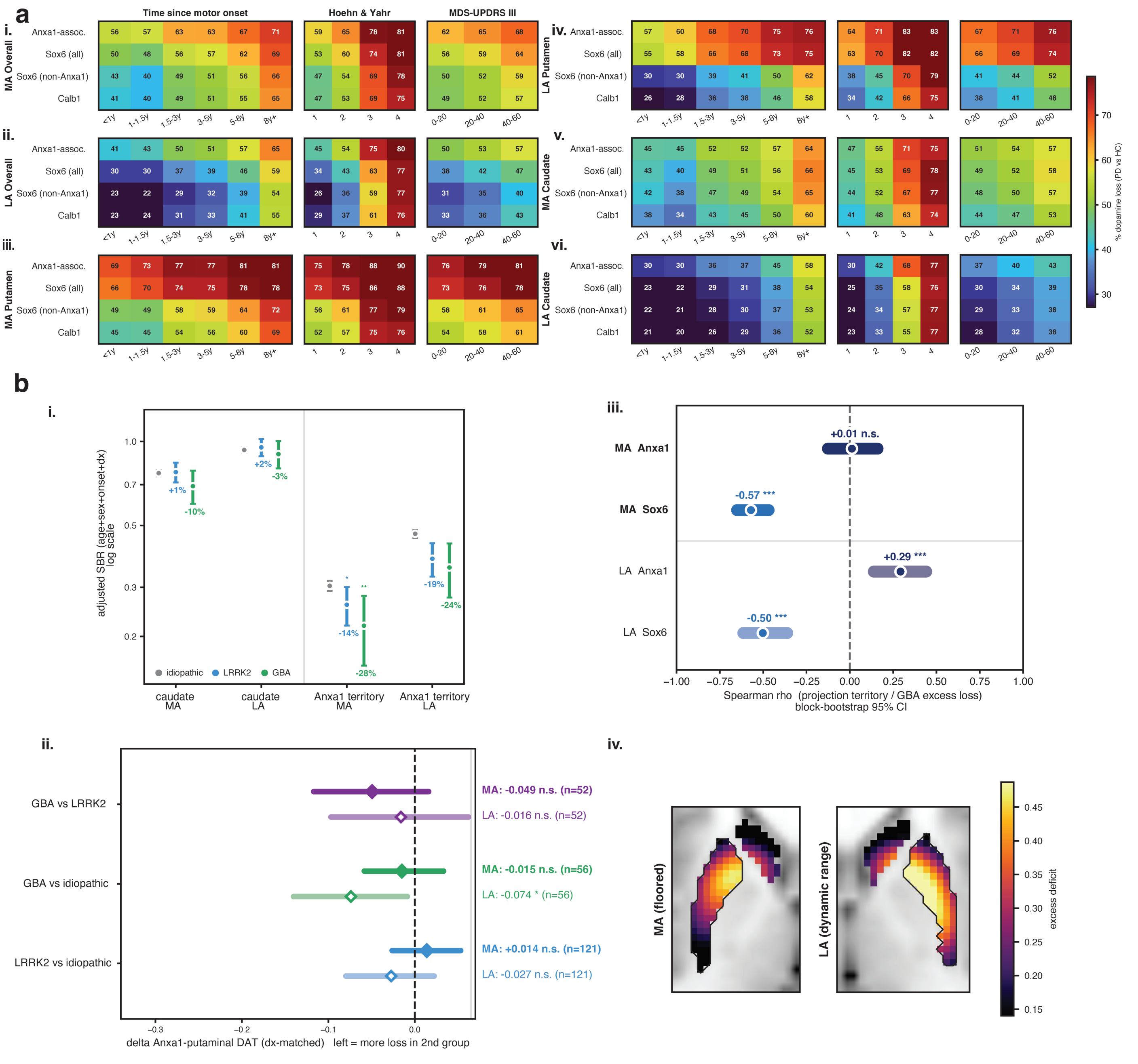

### Supplemental Figure 6

#### Figure 6 Supplementary

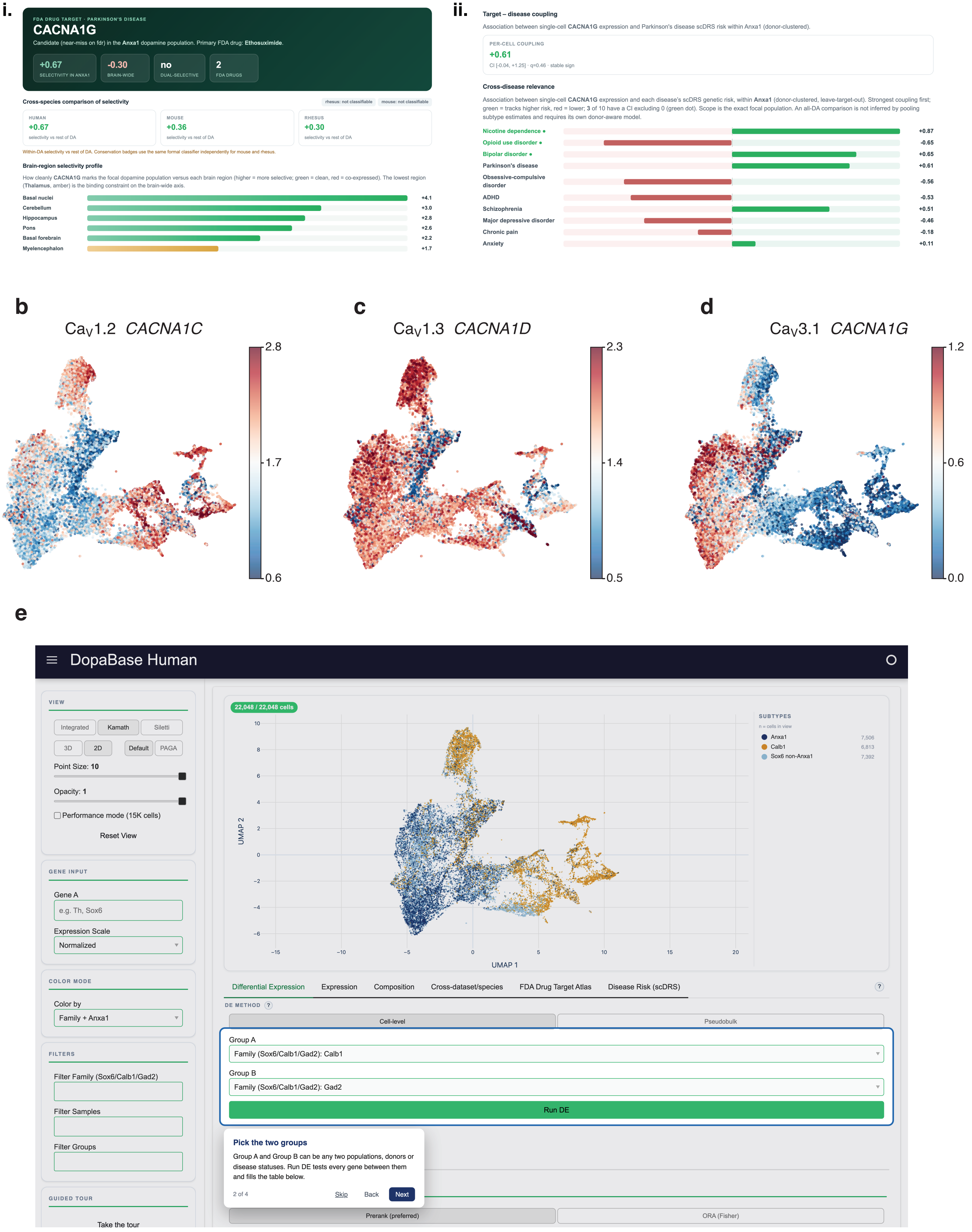
