## Supplemental Figure 2 for "Identification of a human Anxa1-associated dopaminergic population linked to Parkinson’s disease vulnerability and sensorimotor putaminal networks"

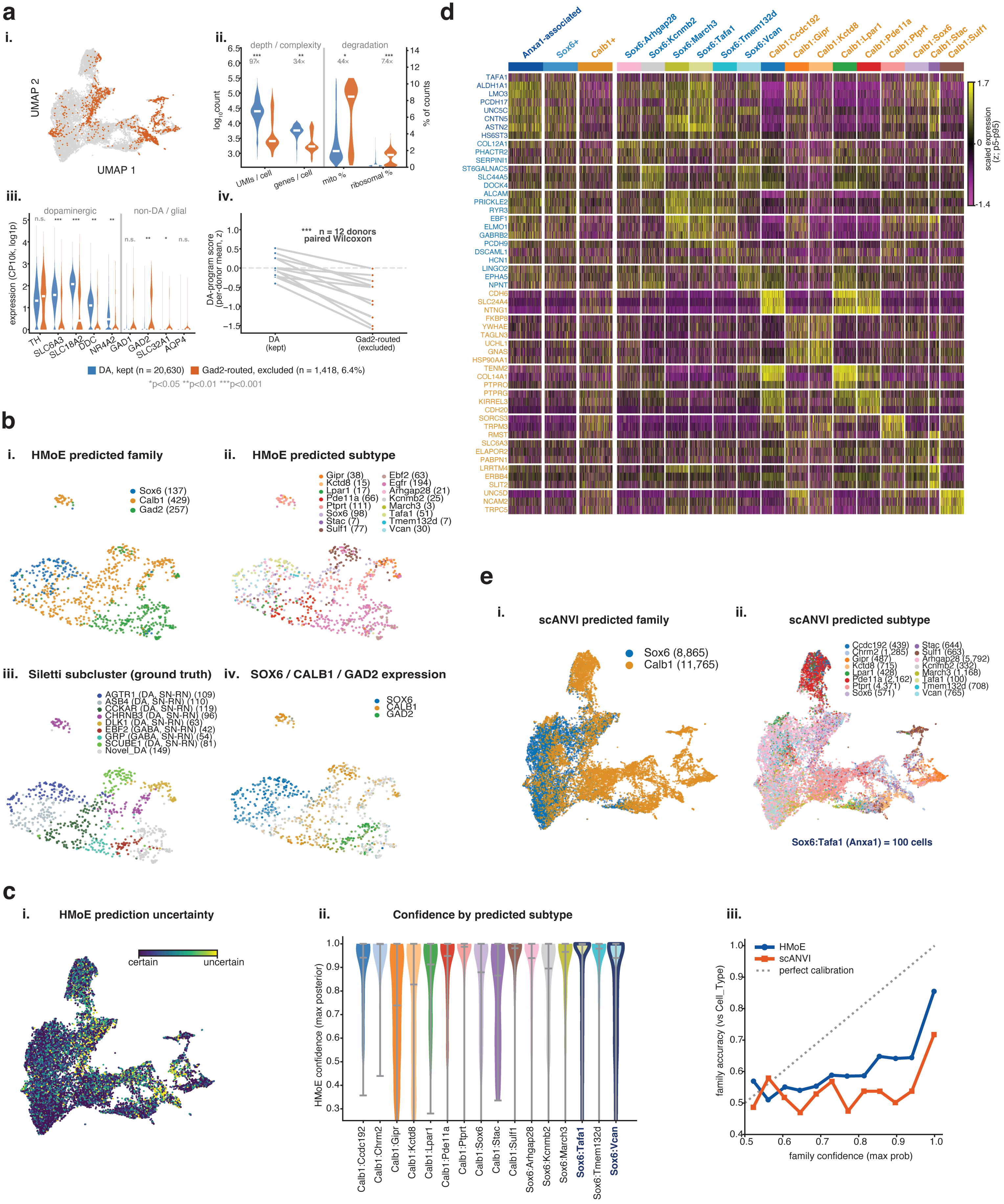

a
i.
UMAP 2
UMAP 1
ii.
depth / complexity
degradation
log10count
% of counts
UMIs / cell
genes / cell
mito %
ribosomal %
97×
34×
44×
74×
iii.
dopaminergic
non-DA / glial
expression (CP10k, log1p)
TH
SLC6A3
SLC18A2
DDC
NR4A2
GAD1
GAD2
SLC32A1
AQP4
iv.
n = 12 donors
paired Wilcoxon
DA-program score (per-donor mean, z)
DA (kept)
Gad2-routed (excluded)
DA, kept (n = 20,630)
Gad2-routed, excluded (n = 1,418, 6.4%)
*p<0.05 **p<0.01 ***p<0.001
b
i. HMoE predicted family
Sox6 (137)
Calb1 (429)
Gad2 (257)
ii. HMoE predicted subtype
Gipr (38)
Kctd8 (15)
Lpar1 (17)
Pde11a (66)
Ptprt (111)
Sox6 (98)
Stac (7)
Sulf1 (77)
Ebf2 (63)
Egfr (194)
Arhgap28 (21)
Kcnmb2 (25)
March3 (3)
Tafa1 (51)
Tmem132d (7)
Vcan (30)
iii. Siletti subcluster (ground truth)
AGTR1 (DA, SN-RN) (109)
ASB4 (DA, SN-RN) (110)
CCKAR (DA, SN-RN) (119)
CHRNB3 (DA, SN-RN) (96)
DLK1 (DA, SN-RN) (63)
EBF2 (GABA, SN-RN) (42)
GRP (GABA, SN-RN) (54)
SCUBE1 (DA, SN-RN) (81)
Novel_DA (149)
iv. SOX6 / CALB1 / GAD2 expression
SOX6
CALB1
GAD2
c
i. HMoE prediction uncertainty
certain
uncertain
ii. Confidence by predicted subtype
HMoE confidence (max posterior)
Calb1:Ccdc192
Calb1:Chrm2
Calb1:Gipr
Calb1:Kctd8
Calb1:Lpar1
Calb1:Pde11a
Calb1:Ptprt
Calb1:Sox6
Calb1:Stac
Calb1:Sulf1
Sox6:Arhgap28
Sox6:Kcnmb2
Sox6:March3
Sox6:Tafa1
Sox6:Tmem132d
Sox6:Vcan
iii.
HMoE
scANVI
perfect calibration
family accuracy (vs Cell_Type)
family confidence (max prob)
d
Anxa1-associated
Sox6+
Calb1+
Sox6:Arhgap28
Sox6:Kcnmb2
Sox6:March3
Sox6:Tafa1
Sox6:Tmem132d
Sox6:Vcan
Calb1:Ccdc192
Calb1:Gipr
Calb1:Kctd8
Calb1:Lpar1
Calb1:Pde11a
Calb1:Ptprt
Calb1:Sox6
Calb1:Stac
Calb1:Sulf1
TAFA1
ALDH1A1
LMO3
PCDH17
UNC5C
CNTN5
ASTN2
HS6ST3
COL12A1
PHACTR2
SERPINI1
ST6GALNAC5
SLC44A5
DOCK4
ALCAM
PRICKLE2
RYR3
EBF1
ELMO1
GABRB2
PCDH9
DSCAML1
HCN1
LINGO2
EPHA5
NPNT
CDH6
SLC24A4
NTNG1
FKBP8
YWHAE
TAGLN3
UCHL1
GNAS
HSP90AA1
TENM2
COL14A1
PTPRO
PTPRG
KIRREL3
CDH20
SORCS3
TRPM3
RMST
SLC6A3
ELAPOR2
PABPN1
LRRTM4
ERBB4
SLIT2
UNC5D
NCAM2
TRPC5
scaled expression (z; p5–p95)
1.7
0
−1.4
e
i. scANVI predicted family
Sox6 (8,865)
Calb1 (11,765)
ii. scANVI predicted subtype
Ccdc192 (439)
Chrm2 (1,285)
Gipr (487)
Kctd8 (715)
Lpar1 (428)
Pde11a (2,162)
Ptprt (4,371)
Sox6 (571)
Stac (644)
Sulf1 (663)
Arhgap28 (5,792)
Kcnmb2 (332)
March3 (1,168)
Tafa1 (100)
Tmem132d (708)
Vcan (765)
Sox6:Tafa1 (Anxa1) = 100 cells
