## Supplemental Figure 5 for "Identification of a human Anxa1-associated dopaminergic population linked to Parkinson’s disease vulnerability and sensorimotor putaminal networks"

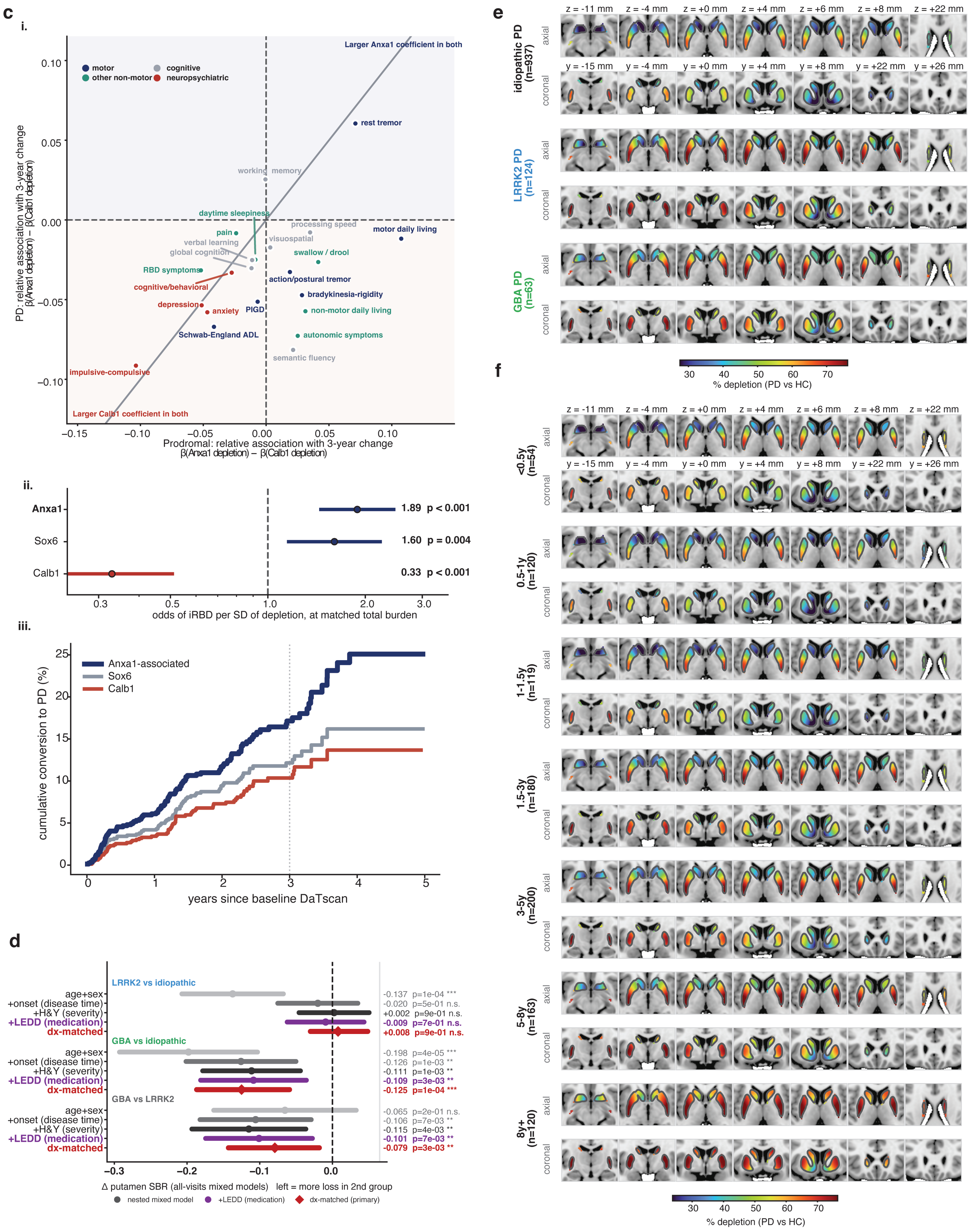
c
i.
Larger Anxa1 coefficient in both
motor
cognitive
other non-motor
neuropsychiatric
rest tremor
working memory
daytime sleepiness
processing speed
motor daily living
pain
verbal learning
visuospatial
global cognition
swallow / drool
RBD symptoms
cognitive/behavioral
action/postural tremor
bradykinesia-rigidity
depression
anxiety
PIGD
non-motor daily living
Schwab-England ADL
autonomic symptoms
semantic fluency
impulsive-compulsive
Larger Calb1 coefficient in both
PD: relative association with 3-year change
β(Anxa1 depletion) − β(Calb1 depletion)
Prodromal: relative association with 3-year change
β(Anxa1 depletion) − β(Calb1 depletion)
0.10
0.05
0.00
−0.05
−0.10
−0.15
−0.10
−0.05
0.00
0.05
0.10
ii.
Anxa1
Sox6
Calb1
1.89 p < 0.001
1.60 p = 0.004
0.33 p < 0.001
0.3
0.5
1.0
2.0
3.0
odds of iRBD per SD of depletion, at matched total burden
iii.
Anxa1-associated
Sox6
Calb1
cumulative conversion to PD (%)
years since baseline DaTscan
0
5
10
15
20
25
0
1
2
3
4
5
d
LRRK2 vs idiopathic
age+sex
+onset (disease time)
+H&Y (severity)
+LEDD (medication)
dx-matched
-0.137 p=1e-04 ***
-0.020 p=5e-01 n.s.
+0.002 p=9e-01 n.s.
-0.009 p=7e-01 n.s.
+0.008 p=9e-01 n.s.
GBA vs idiopathic
age+sex
+onset (disease time)
+H&Y (severity)
+LEDD (medication)
dx-matched
-0.198 p=4e-05 ***
-0.126 p=1e-03 **
-0.111 p=1e-03 **
-0.109 p=3e-03 **
-0.125 p=1e-04 ***
GBA vs LRRK2
age+sex
+onset (disease time)
+H&Y (severity)
+LEDD (medication)
dx-matched
-0.065 p=2e-01 n.s.
-0.106 p=7e-03 **
-0.115 p=4e-03 **
-0.101 p=7e-03 **
-0.079 p=3e-03 **
-0.3
-0.2
-0.1
0.0
Δ putamen SBR (all-visits mixed models) left = more loss in 2nd group
nested mixed model
+LEDD (medication)
dx-matched (primary)
e
z = -11 mm
z = -4 mm
z = +0 mm
z = +4 mm
z = +6 mm
z = +8 mm
z = +22 mm
y = -15 mm
y = -4 mm
y = +0 mm
y = +4 mm
y = +8 mm
y = +22 mm
y = +26 mm
idiopathic PD (n=937)
LRRK2 PD (n=124)
GBA PD (n=63)
axial
coronal
30
40
50
60
70
% depletion (PD vs HC)
f
<0.5y (n=54)
0.5-1y (n=120)
1-1.5y (n=119)
1.5-3y (n=180)
3-5y (n=200)
5-8y (n=163)
8y+ (n=120)
axial
coronal
30
40
50
60
70
% depletion (PD vs HC)
